# The nucleolar protein GNL3 prevents resection of stalled replication forks

**DOI:** 10.1101/2022.10.27.514025

**Authors:** Rana Lebdy, Marine Canut, Julie Patouillard, Jean-Charles Cadoret, Anne Letessier, Josiane Ammar, Jihane Basbous, Serge Urbach, Benoit Miotto, Angelos Constantinou, Raghida Abou Merhi, Cyril Ribeyre

## Abstract

DNA replication requires specific proteins that protect replication forks and so prevent the formation of DNA lesions that may damage the genome. Identification of new proteins involved in these processes is essential to understand how cancer cells tolerate DNA lesions. Here we show that human GNL3/nucleostemin, a GTP-binding protein localized mostly in the nucleolus and highly expressed in cancer cells, prevents nuclease-dependent resection of nascent DNA in response to exogenous replication stress. We demonstrate that inhibition of origin firing decreases this resection, indicating that the increased replication origin firing seen upon GNL3 depletion mainly accounts for the observed DNA resection. We show that GNL3 and DNA replication initiation factor ORC2 interact in the nucleolus and that the concentration of GNL3 in the nucleolus is required to limit DNA resection in response to replicative stress. We propose that the accurate control of origin firing by GNL3, possibly through the regulation of ORC2 sub-nuclear localization, is critical to prevent nascent DNA resection in response to replication stress.

## Introduction

In all cells, DNA replication must occur precisely before their division to ensure faithful transmission of the genome. In humans, accurate DNA replication is particularly important for stem cells and for preventing premature aging and/or cancer (Macheret and Halazonetis, 2015; Schumacher et al., 2021). Replication must occur correctly in space and time to ensure that the whole genome is copied entirely once per cell cycle with no under-replicated or over-replicated regions. Moreover, the replication forks – the sites at which the replication machinery (the replisome) replicates DNA – must be free of impediments that perturb their progression because collapsed replication forks can result in DNA lesions.

DNA replication initiates from specific sites distributed all over the genome, called replication origins (Fragkos et al., 2015; Mechali, 2010). Initiation of replication is a two-step process. First, the origins are ‘licensed’ for replication by binding of the origin recognition complex (ORC, composed of six subunits, ORC1–6) and the replicative helicase MCM2–7, which forms the pre-replicative complex. Second, origin firing (the start of DNA synthesis) which requires activation of cyclin-dependent kinases and CDC7/DBF4 kinases. Although the ORC complex is mainly responsible for initiating DNA replication, it also has other functions. For example, one of the ORC subunits, ORC2, plays roles at centromeres and in sister chromatid cohesion independently of the ORC complex (Bauwens et al., 2021; Huang et al., 2016; MacAlpine et al., 2010; Prasanth et al., 2004; Shimada and Gasser, 2007).

After DNA replication starts, the progression of the replisome may be perturbed by factors of endogenous and exogenous origin that induce replication stress (Lambert and Carr, 2013). The main pathway activated to prevent fork collapse and genomic instability, the ATR–Chk1 checkpoint, prevents further progress through S phase, thus providing time for stalled forks to be stabilized to avoid formation of DNA lesions (Zeman and Cimprich, 2014). Many other proteins, for example BRCA1, protect stalled forks by preventing the action of specific nucleases like MRE11 or CtIP (Berti et al., 2020; Liao et al., 2018; Rickman and Smogorzewska, 2019). ATR–Chk1 also maintains genomic stability by limiting the firing of replication origins in response to replication stress (Blow et al., 2011; Courtot et al., 2018; Toledo et al., 2013). WEE1, a kinase that limits entry into mitosis by inhibiting CDK1, acts in a similar way (Beck et al., 2012; Moiseeva et al., 2019; Toledo et al., 2013).

We previously used the iPOND (isolation of proteins on nascent DNA) method coupled with mass spectrometry (iPOND-MS) to identify novel factors associated with replication forks (Lebdy et al., 2023; Lossaint et al., 2013; Ribeyre et al., 2016). Here, we used an siRNA screen to identify those novel factors whose depletion increases the number of DNA lesions in response to replication stress. The protein whose depletion had the greatest effect was GNL3 (also known as nucleostemin), a GTP-binding protein localized mainly in the nucleolus which is highly expressed in stem cells and cancer cell lines (Tsai and McKay, 2002). Previous studies found that GNL3 depletion leads to activation of the DNA damage response during S phase (Lin et al., 2013; Meng et al., 2013; Yamashita et al., 2013). GNL3 is recruited to DNA double-strand breaks (DSBs), and its depletion prevents RAD51 – a key protein for DSBs repair by homologous recombination – from being recruited at DSBs and hydroxyurea (HU)-induced lesions (Lin et al., 2013; Meng et al., 2013). Consistent with this, GNL3-depleted cells are more sensitive to HU (Lin et al., 2014) and are less able to repair DSBs by homologous recombination (Meng et al., 2013). The current model suggests that GNL3 in the nucleoplasm maintains genome stability in S phase by being recruited to DNA lesions to stabilize RAD51 (Tsai, 2014). The precise functions of GNL3 in S phase, its role in DNA replication and genome stability, are poorly understood, however. In this report, we demonstrate that the concentration of GNL3 in the nucleolus is required to protect stalled replication forks by limiting replication origins firing.

## Results

### GNL3 prevents DNA resection of stalled replication forks

We reported previously our use of the iPOND method to identify novel factors associated with replication forks (Lebdy et al., 2023). Briefly, we pulse-labelled newly synthesized DNA in Hela S3 cells with 5-ethynyl-2’-deoxyuridine (EdU, a nucleoside analogue of thymidine that can be labelled by Click chemistry) or pulsed with EdU then chased for two hours with thymidine, then we purified the proteins associated with EdU. Those proteins that were significantly enriched in the pulse-labelled samples when compared to the chase were defined as components of the replisome (Lebdy et al., 2023). These components included many proteins that were not previously known to be associated with nascent DNA. To select candidates for further analysis, we designed an orthogonal approach based on a mini screen using 25 individual endoribonuclease-prepared siRNAs (esiRNAs; against 24 candidates plus a negative control esiRNA against EGFP). We wished to focus on proteins required to protect DNA integrity, in this case their depletion should increase the number of DNA lesions upon treatment with exogenous molecules that enhance replication stress. We analyzed DNA lesions by quantifying the amount of γH2A.X phosphorylation after 4 hours of replication stress due to treatment with 1 µM camptothecin (CPT, an inhibitor of DNA topoisomerase 1). Briefly, HCT116 cells growing in 96 well plates were transfected with each of the 25 esiRNAs. Forty-eight hours after transfection, the cells were treated for 4 hours with 1 μM CPT and the amount of γH2A.X in the nucleus (seen by staining with DAPI) was analyzed by immunofluorescence microscopy using a Celigo high-throughput microscope (Fig EV1A). We ranked the effects of the 25 esiRNAs based on the amount of γH2A.X and found that GNL3 ranked highest, suggesting that it may be important to tolerate replication stress (Fig EV1B). This is consistent with earlier results showing that GNL3 depletion leads to activation of the DNA damage response during S phase and GNL3-depleted cells are more sensitive than control cells to hydroxyurea (HU), an inducer of replication stress (Lin et al., 2013; Lin et al., 2014; Meng et al., 2013; Yamashita et al., 2013). Since we found more γH2A.X in the nucleus of CPT-treated cells depleted of GNL3 than in control cells (Fig EV1B), we investigated further whether GNL3 regulates replication fork progression in the presence of CPT. To do so we depleted GNL3 (Fig 1A) and labelled cells for 30 min with IdU followed by labelling for 30 min with CldU in the presence or absence of 1 μM CPT and measured the length of both tracks to obtain the CldU/IdU ratio (Fig 1B). As expected, addition of CPT strongly reduced the CldU/IdU ratio, however, depletion of GNL3 had no additional impact (Fig 1B, Fig EV1C). This indicates that GNL3 has no great influence on replication fork progression during brief treatments with CPT. When the cells were treated with CPT for 1, 2 and 4 hours (Fig EV1D), CPT treatment induced rapid phosphorylation of the DNA damage response kinase Chk1 on Ser 345, as expected, however, the kinetics of its phosphorylation was not markedly affected by GNL3 depletion, further supporting our conclusion that GNL3 does not affect fork progression in response to CPT. By contrast, after 4 hours of treatment with CPT the level of phosphorylation of RPA on both Ser 33 and Ser 4/8 was higher in the absence of GNL3 than in the controls (Fig EV1D). To determine if this effect was specific to CPT, we performed the same experiment but treated the cells with HU or etoposide (ETP), a topoisomerase 2 inhibitor. Treatment with 5 mM HU or 10 μM ETP induced phosphorylation of Chk1 on serine 345 in control cells but, as with CPT, no obvious difference was seen when GNL3 was depleted (Fig 1C and Fig EV1E). Also, as with CPT, we observed stronger phosphorylation of RPA on Ser 33 and Ser 4/8 in the absence of GNL3 than in control cells after 4 hours treatment with HU (Fig 1C) and after 2 hours treatment with ETP (Fig EV1E). Thus, we hypothesized that GNL3 depletion may not impact replication stress signaling through Chk1 but, rather, the stability of stalled replication forks, since RPA phosphorylation is a marker of DNA resection (Soniat et al., 2019). Several proteins, including BRCA1, BRCA2 and FANCD2, have been shown to protect nascent DNA from resection in response to replication stress (Rickman and Smogorzewska, 2019). To test if GNL3 protects nascent strand DNA, we sequentially labelled cells with IdU and CldU for 30 min each and then treated the cells with HU for 4 hours (Fig 1D). In the controls, the CldU/IdU ratio was close to 1, indicating that the nascent DNA was protected from extensive degradation, as expected. In cells depleted of GNL3, by contrast, the CldU/IdU ratio was significantly lower (Fig 1D, Fig EV1F), indicating DNA resection at the fork by nuclease(s). Likewise, we saw similar effects in response to CPT (Fig EV1G) and ETP (Fig EV1H), consistent with the increased level of RPA phosphorylation induced by these agents in GNL3-depleted cells.

**Figure 1.**
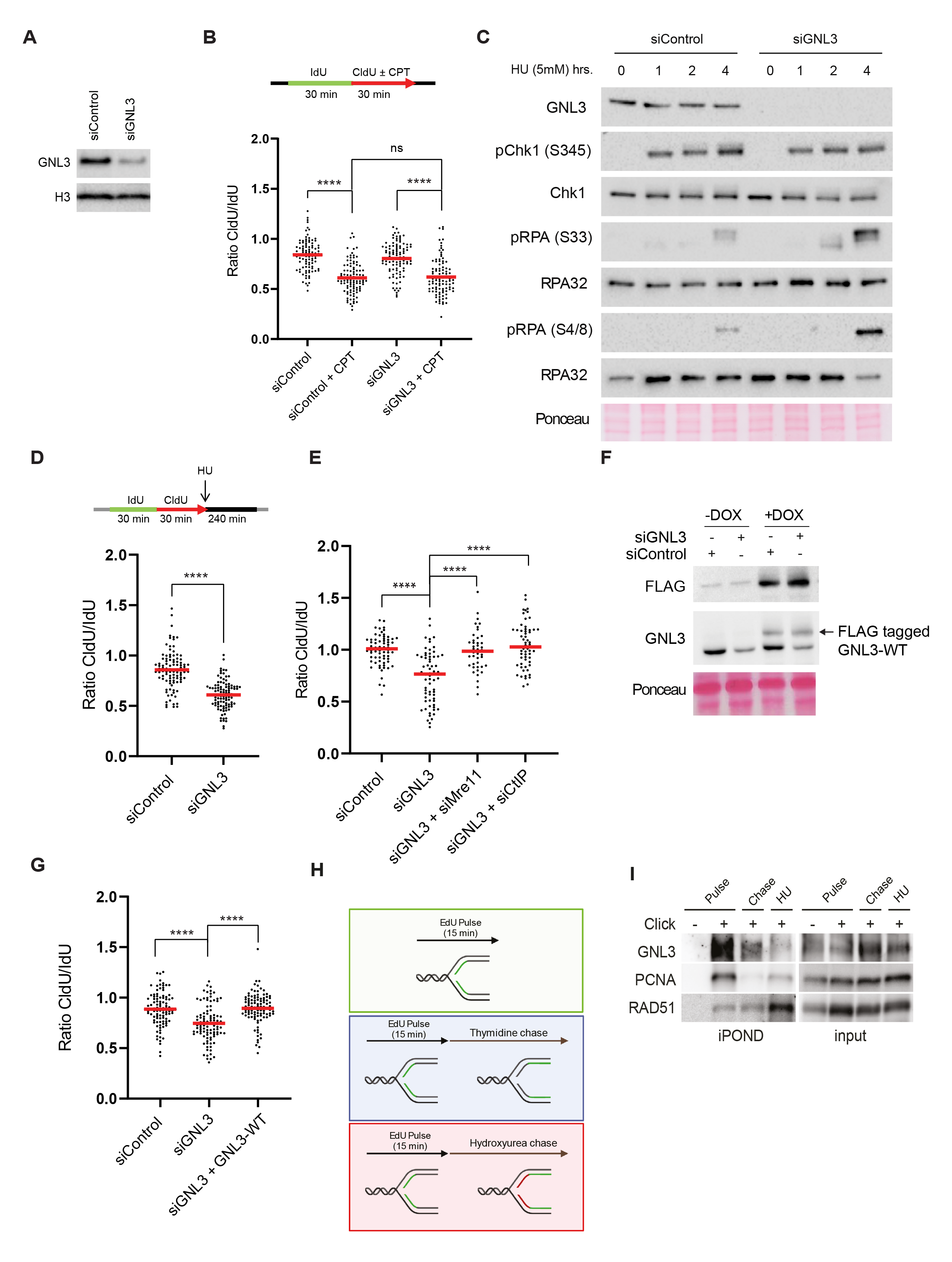
GNL3 prevents DNA resection of stalled replication forks. **A.** Western-blot analysis of HeLa S3 cells depleted with a pool of 4 siRNA targeting GNL3 (siGNL3) or not (siControl). **B.** HeLa S3 cells were sequentially labelled for 30 min with IdU and for 30 min with CldU with or without 1 μM CPT. Ratios between CldU and IdU are plotted, the red line indicates the median. For statistical analysis Mann-Whitney test was used; ****p<0.0001. ns, not significant. **C.** Western-blot analysis of HeLa S3 cells treated with 5 mM HU during the indicated time. **D.** HeLa cells were sequentially labelled for 30 min with IdU and for 30 min with CldU then treated with 5 mM HU for 240 min. The ratio between CldU and IdU is plotted, the red line indicates the median. For statistical analysis Mann-Whitney test was used; ****p<0.0001. **E.** HeLa S3 were sequentially labelled for 30 min with IdU and for 30 min with CldU then treated with 5 mM HU for 240 min. The ratio between CldU and IdU is plotted, the red line indicates the median. For statistical analysis Mann-Whitney test was used; ****p<0.0001. **F.** Western-blot analysis of Flp-in T-Rex HeLa cells expressing GNL3-WT tagged with FLAG. Cells were first transfected with siControl or siGNL3 for 48 hrs then expression of GNL3-WT (resistant to the siRNA against GNL3) was induced using 10 μg/ml of doxycycline (DOX) for 16 hrs. **G.** Flp-in T-Rex HeLa cells were sequentially labelled for 30 min with IdU and for 30 min with CldU then treated with 5 mM HU for 240 min. The ratio between CldU and IdU is plotted, the red line indicates the median. For statistical analysis Mann-Whitney test was used; ****p<0.0001. **H.** Experimental set-up of iPOND experiment. **I.** iPOND experiment analyzed by Western-blot. Cells were pulsed with 15 min EdU and chased for 2 hrs with 10 μM thymidine or 5 mM HU. In no click sample, biotin-TEG azide was replaced by DMSO.

The resection observed in the absence of fork protectors is most probably initiated by the endonuclease activities of MRE11 and CtIP (Rickman and Smogorzewska, 2019). To test further the function of GNL3 as a fork protector, we depleted GNL3 and MRE11, or GNL3 and CtIP, and found that loss of the nucleases prevented the resection seen upon depletion of GNL3 alone (Fig 1E, Fig EV1I, Fig EV1J), further supporting our conclusion that GNL3 protects nascent strand degradation by nucleases. To show definitively that GNL3 protects against DNA resection at stalled replication forks, we depleted the endogenous GNL3 with a specific siRNA and complemented its function by expressing an siRNA-resistant, doxycycline (DOX)-inducible GNL3-FLAG gene in Flp-In T-Rex HeLa cells (Fig 1F). We treated these cells with HU and analyzed the level of resection by IdU and CldU incorporation, as before. Expression of siRNA-resistant GNL3-FLAG suppressed almost completely the increased resection due to GNL3 depletion (Fig 1G, Fig EV1K).

Other proteins known to protect replication forks – BRCA1, RAD51 and FANCD2, for example – accumulate on HU-stalled forks (Dungrawala et al., 2015; Lossaint et al., 2013; Zellweger et al., 2015), suggesting that they may protect them directly from the action of nucleases. To determine whether GNL3 protects stalled replication forks from nucleases in the same way, we used iPOND to identify the proteins on nascent DNA. Cells were pulse labelled for 15 min with EdU and then chased for 2 hours with thymidine or with HU (Fig 1H). As already shown (Dungrawala et al., 2015; Sirbu et al., 2011), treatment with HU increased the recruitment of RAD51 (Fig 1I). By contrast, recruitment of GNL3 was strongly decreased in response to HU, as was PCNA (Figure 4B), indicating that GNL3 does not accumulate at stalled forks. This suggests that the ability of GNL3 to protect from resection may not rely on direct protection from nucleases.

### GNL3 depletion increases the firing of replication origins

To try to understand how GNL3 might protect stalled replication forks from resection we analyzed the impact of GNL3 depletion on DNA replication in basal condition. We found no obvious effect of GNL3 depletion, however, either on the distribution of cells in various phases of the cell cycle whether in an unsynchronized population (Fig EV2A) or in a population synchronized with a thymidine block and released into S-phase (Fig 2A). To confirm this conclusion, we measured the length of S phase by examining the timing of entry into mitosis after a thymidine block, as indicated by phosphorylation of histone H3 on Ser 10 (Prigent and Dimitrov, 2003). Confirming that the length of S phase was unaffected by GNL3 depletion, no sign of early mitotic entry was detected 8 hours after release (Fig EV2B). Ten hours after release, however, we noticed a small increase in the percentage of pH3S10-positive cells in GNL3-depleted cells when compared to the control, suggesting the cells accumulate in mitosis in the absence of GNL3, a phenomenon observed also in breast cancer cells lacking GNL3 (Lin et al., 2014). In those cells, loss of GNL3 increased the number of foci containing the DNA damage response protein 53BP1 (Lin et al., 2014; Yamashita et al., 2013), potentially an indicator of incomplete replication (Harrigan et al., 2011). To test if GNL3 depletion perturb DNA replication, we analyzed its dynamic with DNA combing (Fig 2B): we labelled the cells with IdU for 20 min and then with CldU, for 20 min and observed that GNL3 depletion reduced fork velocity by about 25% (Fig 2C, Fig EV2C). Since the length of S phase is not affected by GNL3 depletion, this may reflect a change in the number of active replication origins. To investigate this possibility, we determined the number of forks per megabase of combed DNA by using a highly accurate assay for global instant fork density (Bialic et al., 2015), which reflects the density of fired replication origins. A significant increase in the number of forks per megabase in GNL3-depleted cells indicated that indeed more origins fire in absence of GNL3 than in control cells (Fig 2D). To confirm this observation, we isolated the chromatin from cells depleted of GNL3 and from control cells and analyzed the presence of markers of origin firing by western blotting. We found more CDC45, MCM2 phosphorylated at Ser 40/41 (pMCM2 S40/41) and PCNA in the chromatin fraction of cells depleted of GNL3 than in control cells (Fig 2E, Fig 2F) confirming that more origins are firing in the absence of GNL3. To investigate whether GNL3 affects the firing of replication origins globally or only at specific regions, as does RIF1 (Yamazaki et al., 2012), we analyzed the effect of GNL3 depletion on replication timing. As expected from previous studies (Cornacchia et al., 2012; Yamazaki et al., 2012), depletion of RIF1 had a substantial impact on replication timing; some regions were delayed and others advanced when compared to the control (Fig EV2D). GNL3 depletion, by contrast, had little or no effect on replication timing (Fig 2G). We conclude that GNL3 depletion increases the firing of replication origins globally without affecting the replication timing.

**Figure 2.**
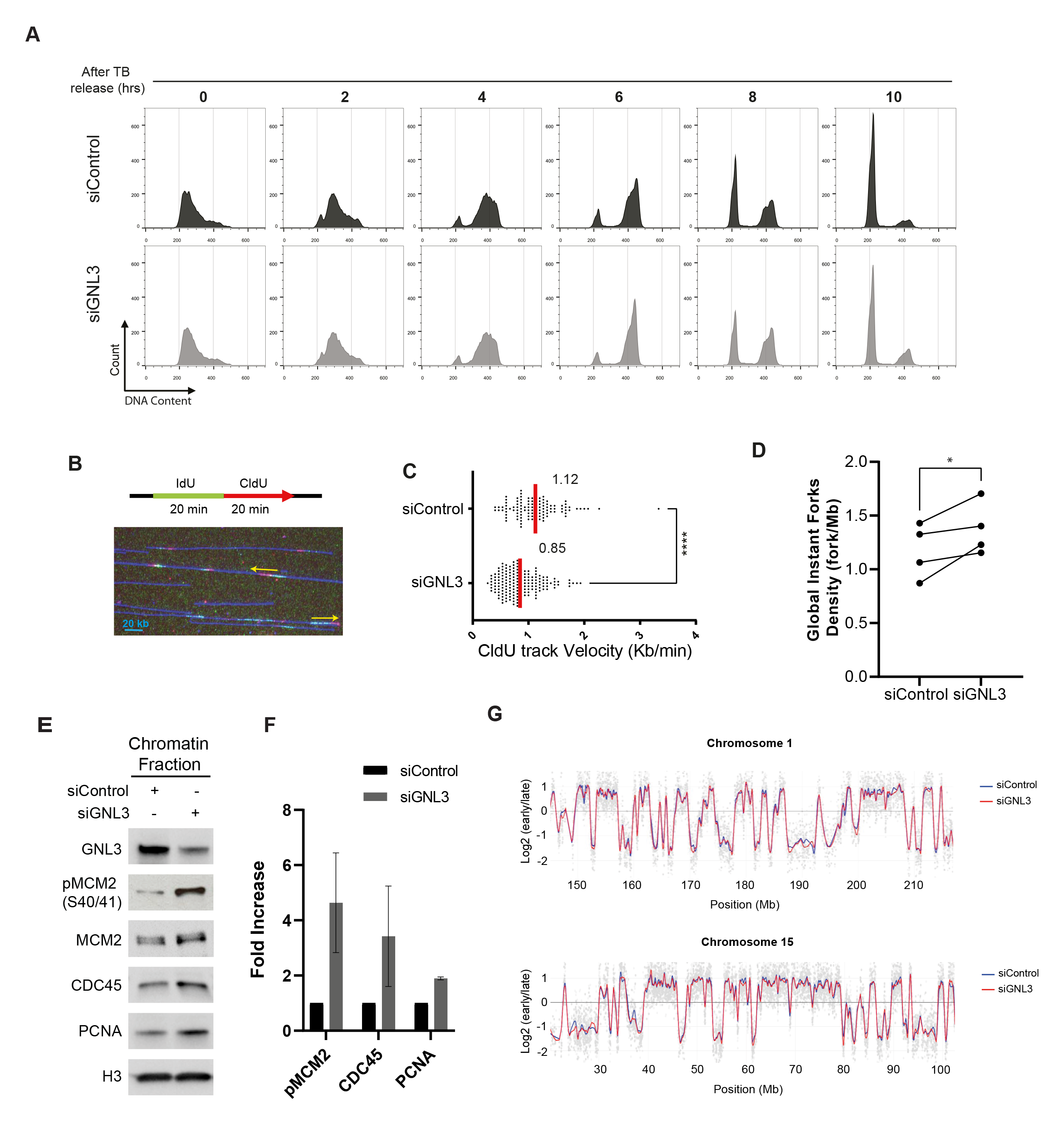
GNL3 depletion increases the firing of replication origins. **A.** Analysis of HeLa S3 cells subjected to thymidine block and released at different timepoints by flow cytometry. DNA was stained with propidium iodide. **B.** DNA combing experiment. HeLa S3 cells were subjected to two consecutive 20 min pulses of IdU and CldU and analyzed by DNA combing. A representative microscopy image of combed DNA molecules containing IdU (red) and CldU (green) tracks in presented with arrows indicating the direction of replication. **C.** Analysis of replication forks velocity by DNA combing. For statistical analysis Mann-Whitney test was used; ****p<0.0001. **D.** Analysis of GIFD (Global Instant Fork Density) by DNA combing in HeLa S3 cells. For statistical analysis paired t-test was used; *p<0.05. **E.** Western-blot analysis of the indicated proteins upon chromatin fractionation. **F.** Quantification of chromatin fractionation based on 3 independent experiments. **G.** Replication timing experiment. HeLa S3 cells were pulse-labelled with BrdU for 90 min and sorted by flow cytometry in two fractions, S1 and S2, corresponding to early and late S-phase. Neo-synthesized DNA was immunoprecipitated with BrdU antibodies. Early and late neo-synthesized DNAs were labeled with Cy3 and Cy5 and hybridized on microarrays. After analyzing with the START-R software, replication-timing profiles can be obtained from two replicates. Shown are the zoomed microarray profiles of the timing of replication on chromosome 1 and chromosome 15 as example. Blue lines represent replication timing from siControl cells and red lines represent siGNL3 cells and grey spots represent the log ratio intensity for each probes of the microarray. Any significantly disturbed regions are detected by START-R software.

### DNA resection in the absence of GNL3 is a consequence of increased origin firing

So far, we show that GNL3 depletion increases replication origin firing and increases DNA resection in response to exogenous inducers of replication stress. Interestingly, the inhibition of WEE1 or ATR increases replication origin firing (Beck et al., 2012; Moiseeva et al., 2017; Moiseeva et al., 2019) and induces DNA lesions in response to HU (Toledo et al., 2013). Importantly this phenotype is partially suppressed by inhibition of origin firing (Toledo et al., 2013), suggesting that increased resection may be a consequence of increased origin firing. We therefore tested the effect of inhibiting ATR or WEE1 on resection in response to HU by sequentially labelling cells with IdU and CldU and then treating them with HU for 4 hours, as before, but in the presence of an inhibitor of ATR or an inhibitor of WEE1 (Fig 3A, Fig EV3A). As predicted, inhibition of ATR (Fig 3B, Fig EV3B) or inhibition of WEE1 (Fig 3C, Fig EV3B) increased resection in response to HU confirming a recent observation for WEE1 inhibition (Elbaek et al., 2022). Moreover, inhibiting the increased origin firing with an inhibitor of CDC7, partially reversed this effect (Fig 3B, Fig EV3B, Fig 3C, FigEV3C). This experiment demonstrates that limiting the number of origins that fire is crucial to preventing resection in response to replication stress. If so, inhibiting origin firing might suppress the HU-induced resection observed upon GNL3 depletion. To test this, we sequentially labelled cells with IdU and CldU for 30 min each and then treated them with HU for 4 hours in the presence of an inhibitor of CDC7 to inhibit replication origin firing. Resection was strongly decreased when CDC7 was inhibited, indicating that in the absence of GNL3 an excess of origin firing in response to HU accounts for the increased resection (Fig 3D, Fig EV3D). Consistent with the decrease in DNA resection, CDC7 inhibition also decreased the phosphorylation of RPA on Ser4/8 (Fig 3E, Fig EV3D).

**Figure 3.**
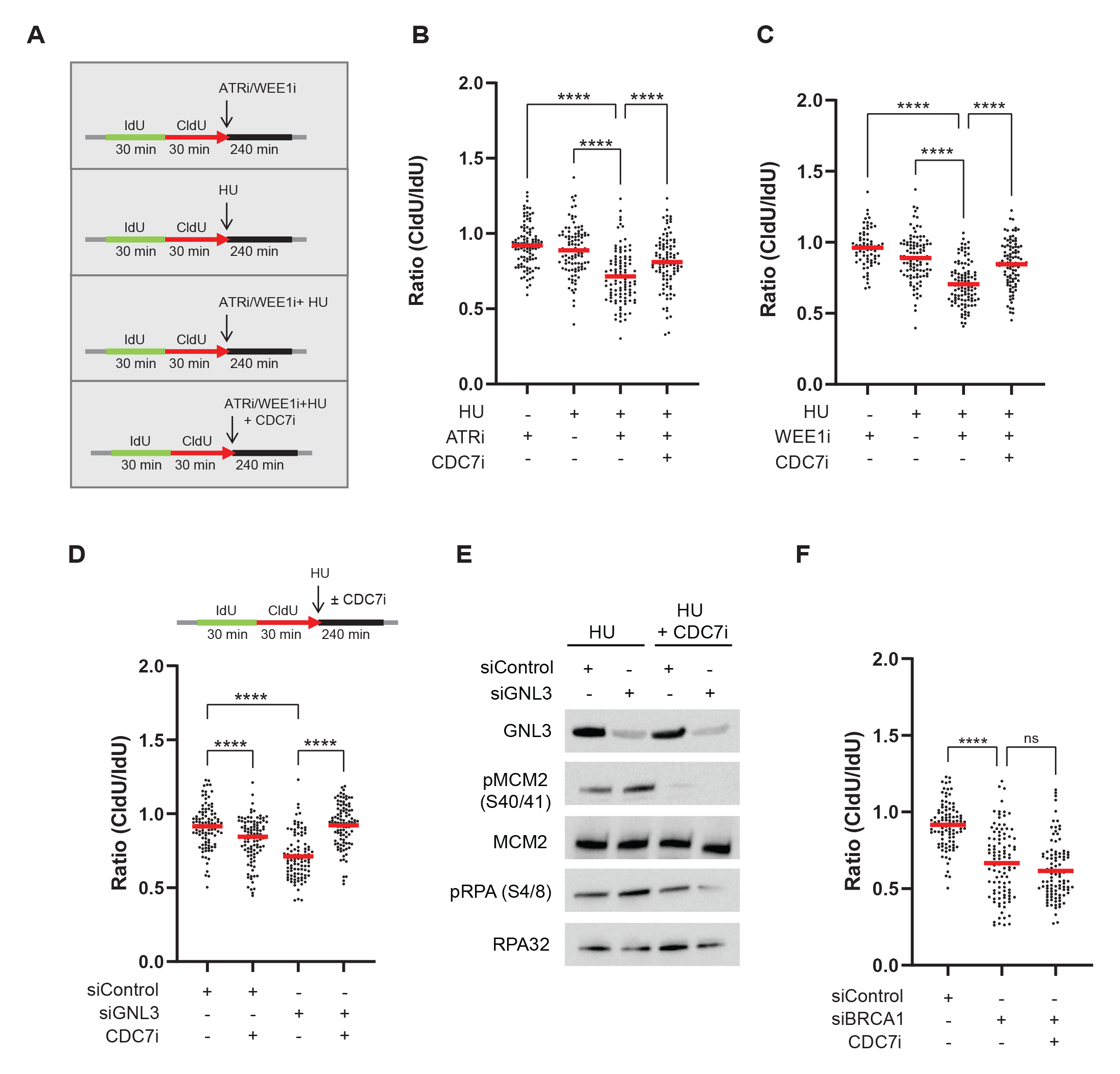
DNA resection in the absence of GNL3 is a consequence of increased origin firing. **A.** HeLa S3 cells were sequentially labelled for 30 min with IdU and for 30 min with CldU then treated or not with 5 mM HU for 240 min with or without 10 μM of ATR VE-821 inhibitor, 500 nM of WEE1 inhibitor AZD1775 or 10 μM of CDC7 inhibitor PHA-767491. **B.** The ratio between CldU and IdU is plotted, the red line indicates the median. For statistical analysis Mann-Whitney test was used; ****p<0.0001. **C.** The ratio between CldU and IdU is plotted, the red line indicates the median. For statistical analysis Mann-Whitney test was used; ****p<0.0001. **D.** HeLa S3 were sequentially labelled for 30 min with IdU and for 30 min with CldU then treated with 5 mM HU for 240 min with or without 10 μM of CDC7 inhibitor PHA-767491. The ratio between CldU and IdU is plotted, the red line indicates the median. For statistical analysis Mann-Whitney test was used; ****p<0.0001. **E.** Western-blot analysis of the indicated proteins upon treatment with 5 mM HU for 240 min with or without 10 μM of CDC7 inhibitor PHA-767491. **F.** HeLa S3 cells were sequentially labelled for 30 min with IdU and for 30 min with CldU then treated with 5 mM HU for 240 min with or without 10 μM of CDC7 inhibitor PHA-767491. The ratio between CldU and IdU is plotted, the red line indicates the median. For statistical analysis Mann-Whitney test was used; ****p<0.0001. ns, not significant.

BRCA1 is recruited to HU-stalled forks (Dungrawala et al., 2015) and its depletion increases DNA resection induced by HU (Schlacher et al., 2012), thus this protein is thought to protect stalled forks from resection by directly blocking nucleases. If this is the case, inhibition of CDC7 should have no effect on protection by BRCA1. To test this prediction, we depleted cells of BRCA1 and measured the level of resection in the absence or presence of the CDC7 inhibitor. As expected, depletion of BRCA1 increased resection; treatment with CDC7 inhibitor, however, did not decrease the level of resection (Fig 3F, Fig EV3F, Fig EV3G), confirming that the resection observed in the absence of BRCA1 is not a consequence of faulty origin firing. Thus, fork protection by BRCA1 differs mechanistically from fork protection by GNL3. We conclude that the enhanced resection observed upon GNL3 depletion is a consequence of increased origin firing.

### GNL3 interacts with ORC2 in the nucleolus

To understand how GNL3 might influence replication origin firing, we used proximity-dependent biotinylation identification (BioID; (Roux et al., 2012)) to identify the proteins in proximity to GNL3 by mass spectrometry. We established a Flp-In T-Rex HEK293 cell line expressing a DOX-inducible GNL3 cDNA fused to the biotin ligase BirA and FLAG. Upon induction with DOX for 16 hours, we observed by immunofluorescence microscopy GNL3-BirA-FLAG mainly in the nucleolus (Fig EV4A). Moreover, by using streptavidin conjugated to Alexa Fluor 488 to detect exogenous biotin, we observed a strong signal (Fig EV4A) demonstrating that GNL3-BirA-FLAG is well localized and can biotinylate proteins in its proximity. In four independent experiments, we induced expression of GNL3-BirA-FLAG with DOX for 16 hours and labelled proteins in its proximity with exogenous biotin for 4 hours. Then we purified the biotinylated proteins on streptavidin beads and analyzed them by mass spectrometry. We calculated the LogRatio of the peptides detected upon addition of DOX and biotin compared to the peptides detected in the negative controls (treatment with either DOX or biotin alone) and represented the data in a Volcano plot (Fig 4A). As expected, GNL3 was highly enriched as well as several nucleolar proteins that are known to be in proximity (e.g., GNL3L, GNL2, DDX21, Ki67 or NPM1). Notably, enrichment of ORC2, one of the components of the origin recognition complex, suggested a possible mechanism in the regulation of replication origin firing by GNL3. To confirm the association of ORC2 with GNL3, we immunoprecipitated each of the proteins and analyzed the immunoprecipitates by western blotting; we found GNL3 in immunoprecipitates of ORC2 and *vice versa* (Fig 4B). Mass spectrometry analysis of the proteins that co-immunoprecipitated when using a specific antibody against ORC2 confirmed the presence of GNL3 and most of the ORC subunits, whereas immunoprecipitation with an irrelevant control IgG contained neither GNL3 nor ORC subunits. Moreover, there was a significant overlap between the co-immunoprecipitated proteins and those found by BioID of GNL3: among the 88 proteins significantly enriched by BioID, 35 were found by coimmunoprecipitation with ORC2 (Fig EV4B) and most of them (24/35) are proteins localized in the nucleolus. This suggests that at least a subset of ORC2 might be localized in the nucleolus and that the interaction between ORC2 and GNL3 is likely to occur in this compartment. The association of GNL3 with chromatin (Fig 2E), however, suggests that GNL3 and ORC2 may also interact at or near replication origins. To test this, we performed GNL3 chromatin immunoprecipitation followed by deep sequencing (ChIP-seq) and found 3412 binding sites for GNL3. We compared these binding sites with ORC2-binding sites (Miotto et al., 2016) but found no significant overlap (Fig 4C, Fig EV4C), indicating that the GNL3–ORC2 interaction may occur mostly in the nucleolus rather than on vicinity of replication origins. To test this, we analyzed the GNL3-ORC2 interaction by using PLA and found most foci at the border of regions that stained lightly with DAPI and that correspond to nucleoli (Fig EV4D), thus supporting our hypothesis. The PLA signal was strongly decreased upon depletion of GNL3, validating its specificity. To validate that the interaction between GNL3 and ORC2 is occurring in the nucleolus, we labelled the nucleolus using an antibody directed against NOP1, a nucleolar protein, before performing PLA between ORC2 and GNL3 (Fig 4D). We could observe a good colocalization between GNL3-ORC2 PLA signal and NOP1, confirming that GNL3 and ORC2 interact mainly in the nucleolus.

**Figure 4.**
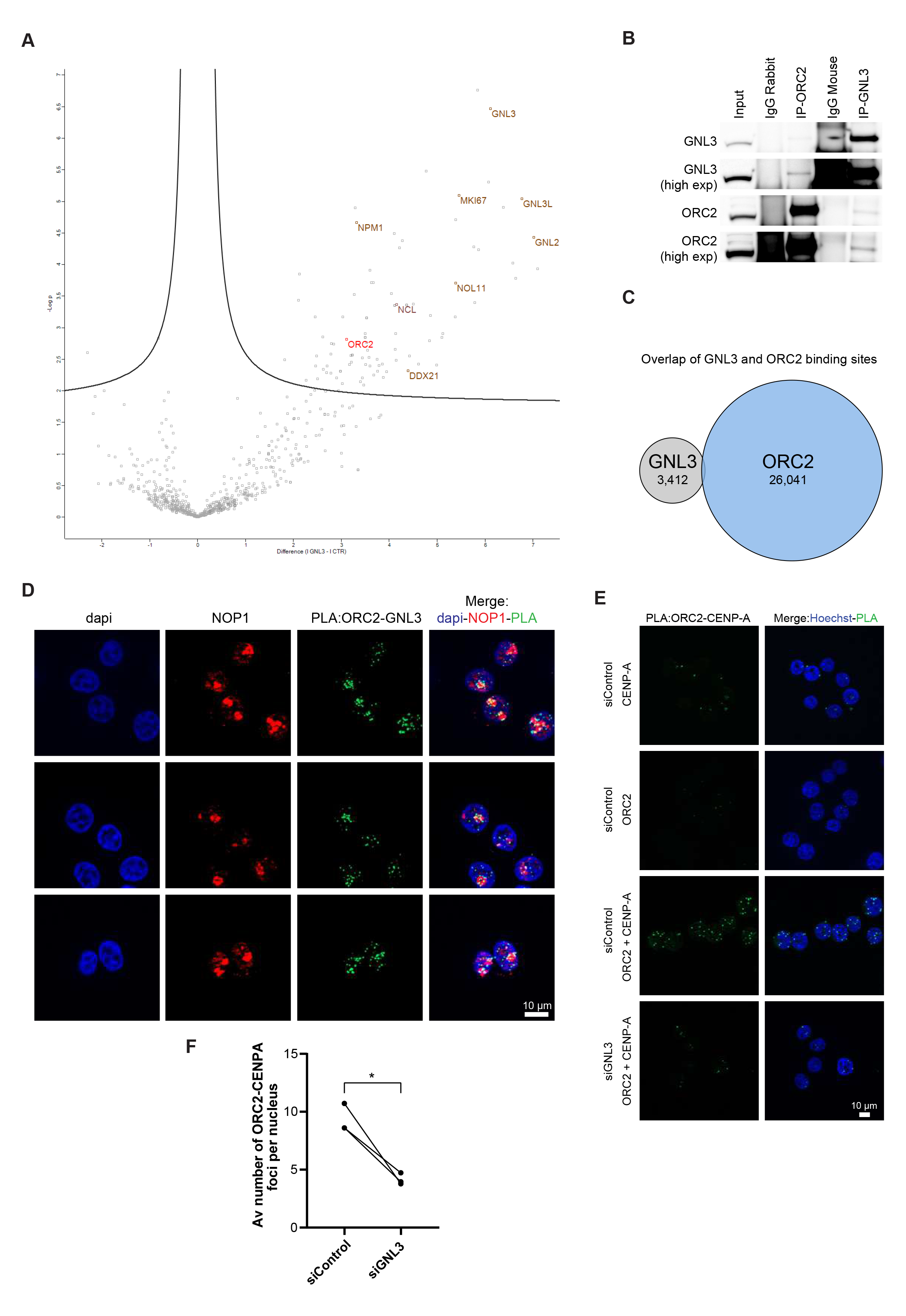
GNL3 interacts with ORC2 in the nucleolus. **A.** GNL3-BioID experiment analyzed by mass spectrometry. Expression of GNL3-BirA-FLAG in HEK293 Flp-in cells was induced with doxycycline for 16 hrs then biotin was added for 4 hrs. For negative controls cells were treated 16 hrs with doxycycline alone or 4 hrs with biotin alone. Each condition was performed four times and analyzed by mass spectrometry. Label free quantification was performed using MaxQuant (Cox and Mann, 2008) and statistical analysis using Perseus (Tyanova et al., 2016). The volcano plot shows the proteins that are significantly enriched upon induction of GNL3-BirA-FLAG and addition of biotin. **B.** Western-blot analysis of GNL3 and ORC2 immunoprecipitates in K562 cells. **C.** Comparison of the genomic location of GNL3 and ORC2. Chromatin immunoprecipitation of GNL3 followed by deep sequencing was performed in HeLa S3. GNL3 binding sites were compared to ORC2 binding sites obtained from Miotto et al. (Miotto et al., 2016). **D.** PLA (proximity ligation assay) analyzing the proximity between ORC2 and GNL3 in HeLa S3 cells that were previously stained with an antibody directed against NOP1. **E.** PLA (proximity ligation assay) analyzing the proximity between ORC2 and CENP-A in HeLa S3 cells using the indicated antibodies. **F.** Graphic representation of the average number of PLA ORC2-CENP-A foci upon 3 independent experiments. For statistical analysis paired t-test was used; *p<0.05.

Our data strongly suggest the presence of a subset of ORC2 into the nucleolus. This is consistent with previous results regarding ORC2 role at centromeres independently of its function in the ORC complex (Bauwens et al., 2021; Huang et al., 2016; Prasanth et al., 2004) since centromeres are often localized in the vicinity of the nucleolus (Padeken et al., 2013; Peng et al., 2023; Wong et al., 2007). In support to this, we observed an interaction between GNL3 and the centromere-specific histone H3 variant CENP-A using PLA (Fig EV4E). We hypothesized that GNL3 may be required for the recruitment of ORC2 at centromeres, a possible readout of its nucleolar localization. To test this, we performed PLA between ORC2 and CENP-A. As expected, many PLA foci of ORC2 and CENP-A were found in normal cells when compared to controls treated with only the antibody against ORC2 or that against CENP-A (Fig 4E). When the cells were depleted of GNL3, however, the average number of PLA foci per cell was reduced by about two-fold (Fig 4F), indicating that ORC2 recruitment at centromeres depends in part on the availability of GNL3. We conclude that the presence of ORC2 at centromeres may reflect its nucleolar localization and could be regulated by GNL3 suggesting that GNL3 may regulate ORC2 sub-nuclear localization.

### The concentration of GNL3 into the nucleolus prevents DNA resection

GNL3 has a long residency time in the nucleolus (Meng et al., 2007). To test if this is important to limit DNA resection and possibly regulates ORC2 sub-nuclear localization, we took advantage of a mutant of GNL3 (GNL3-dB) that is deleted from the B domain (Fig 5A). GNL3-dB has a shorter residency time in the nucleolus and diffuses in the nucleoplasm (Tsai and McKay, 2002, 2005). We depleted endogenous GNL3 with a specific siRNA and expressed siRNA-resistant, DOX-inducible GNL3-dB fused with FLAG in Flp-In T-Rex HeLa cells. GNL3-dB had a level of expression comparable to GNL3-WT (Fig 5B). We next checked GNL3-WT and GNL3-dB localization using immunofluorescence with an antibody directed against FLAG (Fig 5C) and confirmed that deletion of the B domain abolished its specific localization in the nucleolus leading to diffusion in the nucleoplasm (Tsai and McKay, 2002, 2005). To test if a change in the nuclear distribution of GNL3 is affecting its interaction with ORC2 we performed PLA. Consistent with our previous observations (Fig 4D, FigEV4D), we found that exogenous GNL3-WT interacts with ORC2 mostly in proximity of the nucleolus (Fig 5D, FigEV5A). Surprisingly, we observed that the interaction between GNL3-dB and ORC2 was stronger (Fig 5E) and occurs mainly in the nucleoplasm consistent with its localization (Fig EV5B). We conclude that when GNL3 is diffusing in the nucleoplasm it increases its ability to interact with ORC2. Interestingly, by performing immunofluorescence experiment in presence of cytoskeletal buffer (CSK) to remove soluble proteins, we observed that the signal corresponding to GNL3-dB in the nucleoplasm was strongly reduced (Fig 5F). Since ORC2 is mostly associated with chromatin (Ohta et al., 2003), we postulate that GNL3-dB interacts with a fraction of ORC2 that may not be localized to chromatin and could reflect a change in ORC2 sub-nuclear localization. We propose that the inability of GNL3-dB to accumulate in the nucleolus may perturb the sub nuclear localization of ORC2 which in turns may impact the licensing of replication origins. To test if this is causing DNA resection in response to HU, we analyzed the level of resection upon expression of GNL3-dB. As already shown (Fig 1G), expression of GNL3-WT almost completely suppressed the increased resection induced by GNL3 depletion (Fig 5G, Fig EV5C). In contrast, GNL3-dB, did not fully complement the increased resection due to GNL3 depletion (Fig 5G, Fig EV5C). We conclude that diffusion of GNL3 in the nucleoplasm induces DNA resection. To confirm this result, we modulated GNL3-dB expression using different concentrations of DOX (Fig EV5D) and observed that the diffusion in the nucleoplasm was dependent on DOX concentration (Fig EV5E). Strikingly, we observed that the amount of resection was largely correlated with the level of expression of GNL3-dB (Fig 5HC, Fig EV5F), confirming that diffusion of GNL3 in the nucleoplasm leads to DNA resection. We conclude that nucleolar localization of GNL3 is required to prevent excessive DNA resection in response to exogenous replication stress by possibly ensuring the correct sub-nuclear localization of specific proteins such as ORC2.

**Figure 5.**
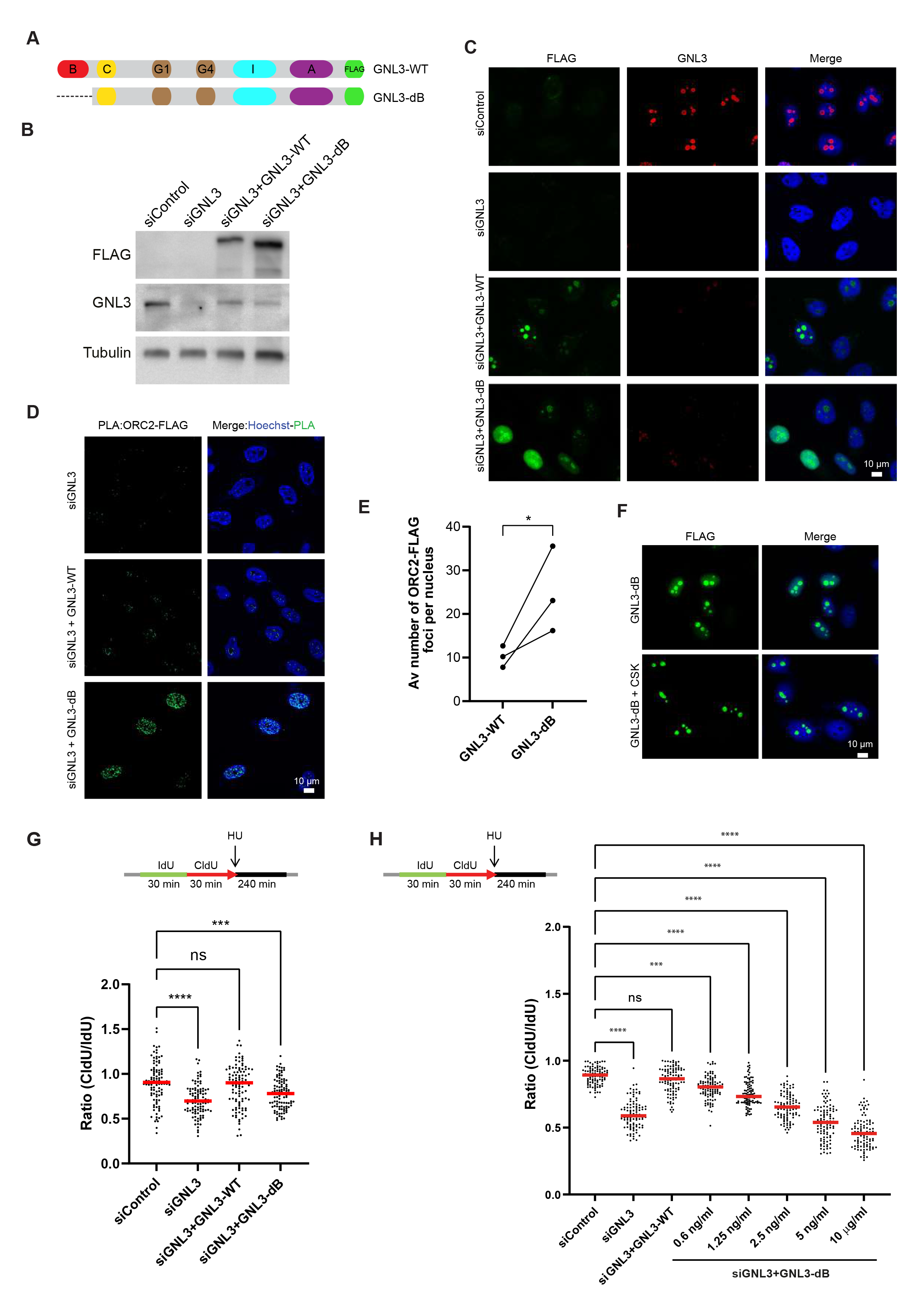
GNL3 accumulation into the nucleolus prevents DNA resection. **A.** Schematic representation of human GNL3 protein with its associated domains (B: basic domain; C: coiled-coil domain; G1: GTP-binding motif 1; G4: GTP-binding motif 4; I: intermediate domain; A: acidic domain). GNL3-WT and GNL3-dB are fused with FLAG. **B.** Western-blot analysis of Flp-In T-Rex HeLa cells expressing exogenous GNL3-WT or GNL3-dB. Cells were transfected with siControl or siGNL3 for 48 hrs then expression of exogenous GNL3-FLAG (resistant to the siRNA against GNL3) was induced using 10 μg/ml of doxycycline for 16 hrs. **C.** Immunofluorescence analysis of Flp-in T-Rex HeLa cells expressing exogenous GNL3-WT or GNL3-dB. **D.** PLA (proximity ligation assay) analyzing the proximity between ORC2 and GNL3-FLAG or GNL3-dB-FLAG in HeLa Flp-In cells upon doxycycline induction. **E.** Graphic representation of the average number of PLA ORC2-FLAG foci upon 3 independent experiments. For statistical analysis paired t-test was used; *p<0.05. **F.** Immunofluorescence experiment of HeLa Flp-In cells expressing GNL3-dB with or without pre-extraction with cytoskeletal buffer (CSK). **G.** Flp-in T-Rex HeLa cells were sequentially labelled for 30 min with IdU and for 30 min with CldU then treated with 5 mM HU for 240 min. The ratio between CldU and IdU is plotted, the red line indicates the median. For statistical analysis Mann-Whitney test was used; ****p<0.0001; ***p<0.001; ns non significant. **H.** Flp-in T-Rex HeLa cells were sequentially labelled for 30 min with IdU and for 30 min with CldU then treated with 5 mM HU for 240 min. The ratio between CldU and IdU is plotted, the red line indicates the median. For statistical analysis Mann-Whitney test was used; ****p<0.0001; ***p<0.001; ns non-significant.

## Discussion

GNL3/nucleostemin was discovered more than twenty years ago as a nucleolar protein required for cell proliferation (Tsai and McKay, 2002) and several studies have highlighted its role(s) in maintaining genome integrity (Tsai, 2014). Here, we investigate the role of GNL3 in response to exogenous replication stress. We demonstrate that GNL3 protects stalled replication forks from resection by endonucleases and this protection depends on the number of replication origins that fire. We show that the long residency time of GNL3 in the nucleolus is required to protect stalled replication from resection by possibly regulating the sub nuclear localization of ORC2. We propose a model in which an excess of fired replication origins induces DNA resection upon treatment with exogenous replication stress (Figure 6).

**Figure 6.**
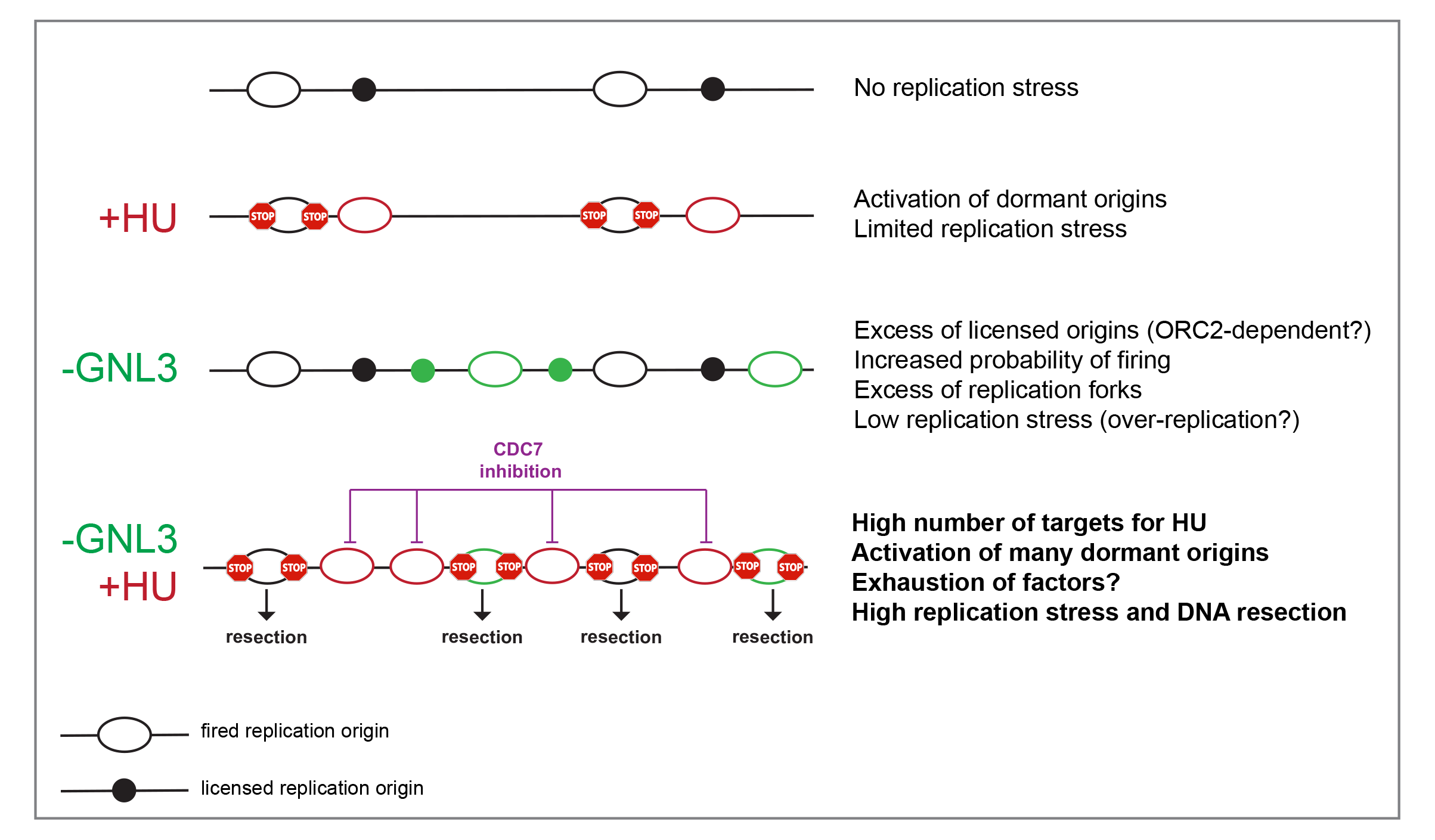
Model. HU treatment stalls active replication forks and activate dormant origins. When GNL3 is impaired, more origins may be licensed increasing the probability of replication origins firing. This could lead to over replication and replicative stress. When GNL3 deficient cells are treated with HU the number of targets for HU is increased leading to activation of more dormant origins, a situation that may lead to an exhaustion of specific protection factors (e.g. BRCA1, RAD51, FANCD2 etc.) that could explain the resection of nascent DNA. Inhibition of origin firing with CDC7 inhibitor counteracts this effect by limiting the need for proteins required for protection.

### GNL3 limits the firing of replication origins

GNL3 depletion induces activation of the DNA damage response during S phase with γH2A.X phosphorylation (Lin et al., 2014; Meng et al., 2013) indicating a strong level of replication stress. Consistent with this and as previously observed (Lin et al., 2014), we noticed a weak enrichment in cells in G2/M cells (Fig 2A, Fig EV2B) that may account for incomplete replication due to replication stress (Harrigan et al., 2011). However, the reason for the endogenous replication stress upon GNL3 depletion was largely unknown. Here we propose that the increase replication origins firing caused by GNL3 depletion may be the major cause of endogenous replicative stress. The regulation of replication origins firing may be less constrained in absence of GNL3 leading to regions under- or over-replicated (Blow et al., 2011) that may induce the accumulation of cells in G2/M. This is supported by the fact that replication timing is not affected upon GNL3 depletion, a measurement that does not represent stochastic variations between individual cells. In addition, recent data suggest that the firing of replication origins is more stochastic than previously thought (Klein et al., 2021; Wang et al., 2021). In addition, the reduced fork velocity observed upon GNL3 depletion may also impair the replication of specific regions. We detected GNL3 in vicinity of replication forks using iPOND coupled with mass spectrometry (Lebdy et al., 2023) and Western-blot (Fig 1F) suggesting that its depletion may directly impairs fork speed. Yet, the recruitment of GNL3 at replication forks is low and it was not detected by other iPOND screens performed in basal conditions (Lopez-Contreras et al., 2013; Sirbu et al., 2013). In fact GNL3 was found at nascent DNA only in FANCJ-knockout cells that tend to heterochromatinize their genome (Peng et al., 2018; Schwab et al., 2013), suggesting that GNL3 may be in proximity of nascent DNA only in specific regions. Interestingly GNL3 is localized in vicinity of centromeres and in proximity of nucleolar associated domains (NADs) that are both heterochromatic regions (Bersaglieri et al., 2022; Peng et al., 2023). Therefore, the weak association of GNL3 to nascent DNA may simply reflect its localization in proximity of heterochromatic regions undergoing DNA replication. This is supported by our recent findings showing that iPOND efficacy is biased by genome organization (Lebdy et al., 2023). Based on this we propose that the slow replication fork speed observed upon GNL3 depletion is more likely a compensation mechanism of the increased replication origin firing (Ge et al., 2007; Ibarra et al., 2008) rather than a direct impact on replication forks speed. The abundance of GNL3 in the nucleolus as well as its interaction with ORC2 (see below) strongly supports this model.

### Excessive replication origin firing induces DNA resection in response to replication stress

We show that GNL3 protects nascent DNA at stalled replication forks from resection by endonucleases. The increased resection seen upon GNL3 depletion, we conclude, is related to the increased replication origin firing because it is suppressed by an inhibitor of CDC7 that decreases origin firing. This conclusion is consistent with data showing that CDC7 inhibition prevents nascent strand resection (Jones et al., 2021; Sasi et al., 2018). We also found that inhibition of ATR or WEE1, both of which increase origin firing (Beck et al., 2012; Moiseeva et al., 2017; Moiseeva et al., 2019), increases the resection of nascent DNA in a CDC7-dependent manner. Collectively, these data suggest that when cells with an excess of fired replication origins are challenged with replication stress, it induces nascent DNA resection. Having more replication origins that fire means more replication forks that may be targeted by exogenous replication stress like HU, which in turn activate more dormant origins (Figure 6). This situation may lead to the exhaustion of specific factors such as RPA causing resection as previously proposed for the occurrence of DNA lesions in the case of ATR or WEE1 inhibition (Toledo et al., 2013). However, we could not detect any sign of RPA exhaustion (data not shown) upon GNL3 depletion. This signifies that RPA exhaustion may not be systematically required for DNA resection and this may rely on the exhaustion of other proteins known for protection such as BRCA1, RAD51 or FANCD2 (Hashimoto et al., 2010; Schlacher et al., 2011; Schlacher et al., 2012). Interestingly, GNL3 is required for RAD51 recruitment upon 24 hours treatment with HU (Meng et al., 2013), yet we could not detect any impact of RAD51 recruitment on chromatin upon GNL3 depletion (data not shown). This suggests that factors that remain to be determined are required to protect stalled replication forks when the regulation of replication origins firing is perturbed.

DNA resection may be largely dependent of replication origin firing. However, the nascent DNA resection that occurs in the absence of BRCA1 was not suppressed by CDC7 inhibition. This confirms a direct role for BRCA1 in protecting nascent DNA. Thus, we conclude that nascent DNA resection can be promoted either by loss of a protein that protects the DNA directly, like BRCA1, or by loss of proteins such as GNL3 or WEE1 that act indirectly. BRCA1, FANCD2 and RAD51 were the first proteins found to act as fork protectors (Hashimoto et al., 2010; Schlacher et al., 2011; Schlacher et al., 2012). Since then, several other proteins have been found to protect stalled forks from resection by nucleases (Berti et al., 2020; Liao et al., 2018; Rickman and Smogorzewska, 2019). Given our findings here, it would be interesting to investigate whether these proteins protect replication forks directly or indirectly.

### Nucleolar concentration of GNL3 is required to prevent DNA resection

To test if GNL3 localization has a role in preventing DNA resection, we expressed a mutant of GNL3, GNL3-dB, that is diffusing in the nucleoplasm (Tsai and McKay, 2002, 2005). We conclude that the concentration of GNL3 in the nucleolus prevents DNA resection in response to replicative stress. One of the main features of GNL3 is its strong residency time in the nucleolus (Meng et al., 2007), this ability may allow GNL3 to inhibit proteins involved in DNA resection by nucleolar sequestration for instance (Wang et al., 2019). To this purpose we screened for proteins localized in proximity of GNL3 using Bio-ID. In addition to known nucleolar proteins our attention was immediately caught by ORC2, one of the best hits. ORC2 is one of the components of the origin recognition complex and therefore is required for the licensing of replication origins. Therefore, in absence of ORC2 less replication origins fire, consequently the inter-origin distance is increased, and fork speed is increased (Shibata et al., 2016). We hypothesized that GNL3 may be able to regulate ORC2 sub-nuclear distribution to limit licensing, something that may explain the increase in replication origins firing we observed upon GNL3 depletion. GNL3 could also regulate ORC2 functions that are not directly linked with the ORC complex but that may influence licensing such as its role at centromeres (Bauwens et al., 2021; Huang et al., 2016; Prasanth et al., 2004), at nuclear pores (Richards et al., 2022) or in sister chromatid cohesion (MacAlpine et al., 2010; Shimada and Gasser, 2007). In support to this, we found that GNL3 depletion decreases ORC2 recruitment to centromeres. ORC2 SUMOylation is required to prevent re-replication of pericentromeric heterochromatin (Huang et al., 2016), suggesting that GNL3 depletion may impair centromeres replication through ORC2 regulation. Since centromeres and replication origins may derives from a common ancestor (Hu and Stillman, 2023), it may also be possible that ORC2 localized at centromeres has a broader role on the control of DNA replication. We have found that GNL3-dB interacts more with ORC2 than GNL3-WT and that this interaction occurs mostly in the nucleoplasm. GNL3-dB may interact with ORC2 from the ORC complex localized on chromatin, however since GNL3-dB is not associated with chromatin in the nucleoplasm we do not favor this possibility. We rather envision that the interaction between ORC2 and GNL3-dB in the nucleoplasm may be due to a release of ORC2 from the nucleolus of from the chromatin. This is supported by the fact that the level of resection increases with the level of GNL3-dB expression, something that could gradually increase the release of ORC2 from the nucleolus. Based on our findings, one possible model is the sequestration of ORC2 by GNL3 that prevents excessive licensing, leading to DNA resection in response to replication stress. This may explain why both GNL3 depletion and expression of GNL3-dB lead to nascent strand resection, since in both cases ORC2 may not be properly sequestrated in the nucleolus. More work is obviously required to demonstrate this model by analyzing for example the distribution of ORC2 on chromatin upon GNL3 loss at genome-wide level.

The ability of GNL3 to prevent excessive firing of replication origins may also be caused to a more global role of GNL3 on nuclear organization possibly related with compartmentation. Although GNL3 is found only in chordates, it belongs to the family of YlqF-related GTPases that is conserved in Eukarya, Bacteria and Archea (Mier et al., 2017; Quiroga-Artigas et al., 2022; Reynaud et al., 2005). GNL3 is the more recent member of the family and seemed to have co-evolved with sub compartments of the nucleolus. Growing evidence indicates that the nucleolus is involved in the 3D organization of the genome (Iarovaia et al., 2019) and particularly of centromeric DNA and heterochromatin (Bersaglieri et al., 2022; Padeken et al., 2013; Peng et al., 2023; Wong et al., 2007). Therefore, GNL3 may play a larger role in the organization of centromeres, or other regions of heterochromatin, by keeping them in proximity to the nucleolus thanks to its long residency time (Meng et al., 2006). For example, GNL3 may mediate interactions between nucleolus and heterochromatin as proposed for Ki-67 (Sobecki et al., 2016; van Schaik et al., 2022) and NPM1 (Holmberg Olausson et al., 2014), two proteins localized to the nucleolar rim like GNL3 (Stenstrom et al., 2020). It has been recently shown that genome organization is a determinant of the locations of replication origins (Emerson et al., 2022), therefore GNL3 may have a broader in role in the regulation of replication origins firing.

## Supporting information

Expanded View Figure 1

Expanded View Figure 2

Expanded View Figure 3

Expanded View Figure 4

Expanded View Figure 5

## Acknowledgments

We thank all the present and former lab members for comments and suggestions on the project and the manuscript. We are grateful to Pierre-Henry Gaillard, Jean-Hugues Guervilly, Maud de Dieuleveult, Anne-Claude Gingras, Yea-Lih Lin and Montpellier Genomic Collection for reagents. We thank Armelle Lengronne, Antoine Aze, Domenico Maiorano, Eric Julien, Sébastien Britton, Olivier Ganier and Joelle Nassar for discussions and comments as well as Marie-Pierre Blanchard and Amélie Sarazzin from the Montpellier Imaging Platform for their support. We acknowledge Carol Featherstone of Plume Scientific Communication Services for professional scientific editing during the preparation of the manuscript. We are grateful to Montpellier Combing Facility (Etienne Schwob and Marjorie Drac) and Montpellier GenomiX facility (Hugues Paranello). Mass spectrometry experiments were carried out using the facilities of the Montpellier Proteomics Platform (PPM, BioCampus). We thank Céline Gongora, Nadia Vie and Naoill Abdellaoui for their help with the use of the Celligo. We thank the 3P5 proteom’IC facility (Johanna Bruce, Cedric Broussard and François Guillonneau) at Institut Cochin, which is supported by the DIM Thérapie Génique Région Ile-de-France, IBiSA, and the Labex GR-Ex. This work was supported by a *Jeunes Chercheurs Jeunes Chercheuses* grant from the *Agence National de la Recherche* (REPLIBLOCK ANR-17-CE12-0034-01), and an Emergence grant from *Cancéropole Grand Sud-Ouest* to Cyril Ribeyre as well as a grant from *Programme labellisé Fondation ARC* to Angelos Constantinou. Rana Lebdy was funded by fellowships from Azm & Saade Association and *Fondation ARC pour la recherche sur le cancer*. Benoit Miotto and Anne Letessier are partners of Labex ‘Who am I?’ (ANR-11-LABX-0071 and ANR-11-IDEX-005-02) and are supported by *Fondation pour la Recherche Medicale* (AJE20151234749), INSERM, CNRS and University Paris Cité. Jean-Charles Cadoret thanks the IdEx Université Paris Cité (ANR-18-IDEX-0001), and the generous legacy from Ms. Suzanne Larzat.

## Author contributions

Conceptualization, R.L. and C.R.; Methodology R.L., M.C., J.P., S.U., J.A., A.L., J-C.C. and C.R.; Validation R.L. and C.R.; Formal Analysis R.L., M.C., J-C.C. and C.R. Funding Acquisition R.L., A.C., R.A.M. and C.R. Supervision R.L., R.A.M. and C.R., Investigation R.L., M.C., J-C.C., A.L. and C.R.; Visualization R.L.; Writing – Original Draft R.L. and C.R.; Writing – Review & Editing R.L., J-C.C., A.L., B.M., J.B., A.C., R.A.M. and C.R. Data Curation R.L., M.C., J-C.C., A.L. and C.R.

## Methods

### Cell lines

HeLa S3 (obtained from ATCC), Flp-In T-Rex 293 (obtained from ThermoFisher) and HeLa Flp-In T-Rex (authenticated with Eurofins, gift from Jean-Hugues Guervilly and Pierre-Henri Gaillard, Centre de Recherche en Cancérologie de Marseille, France) cells were cultured in Dulbecco’s modified Eagle’s media (DMEM). HCT116 (obtained from SIRIC Montpellier Cancer) and K562 (authenticated with Eurofins) cells were cultured in Roswell Park Memorial Institute medium (RPMI). Culture media was supplemented with 10% fetal bovine serum (Biowest) and penicillin/streptomycin (Sigma-Aldrich). Cells were incubated in a 5% CO_2_ at 37^⁰^C. Selection of integrated clones in Flp-In cells were done using hygromycin and blasticidin.

### Inhibitors, drugs and antibiotics

The following reagents were used: etoposide (Sigma-Aldrich E1383), camptothecin (Sigma-Aldrich C9911), hydroxyurea (Sigma-Aldrich H8627), doxycycline (Clontech 631311), hygromycin B Gold (InvioGen), zeocin (Invitrogen 46-0509), blasticidin (InvivoGen), ATR inhibitor VE-821 (TINIB-TOOLS), WEE1 inhibitor AZD1775 (Selleckchem), CDC7 inhibitor PHA-767491 (Selleckchem).

### Plasmids construction

GNL3 cDNA cloned in pDONR223 (obtained from Montpellier Genomic Collection) was introduced using Gateway method in pDEST-pcDNA5-FLAG C-term and pDEST-pcDNA5-BirA-FLAG C-term (gifts from Anne-Claude Gingras, Lunenfeld-Tanenbaum Research Institute at Mount Sinai Hospital, Toronto, Canada). GNL3 truncations were created by gene synthesis (Invitrogen GeneArt Gene Synthesis) and introduced using Gateway method into pDEST-pcDNA5-FLAG C-term.

### Gene silencing

For GNL3 depletion siGENOME SMARTpool (M-016319-00) and individual siRNA oligonucleotide (D-016319-01-0002) were purchased from Dharmacon and transfected using INTERFERin (Polypus transfection). siRNAs against MRE11 and CtIP were provided by Yea-Li Lin (Institut de Génétique Humaine, Montpellier) and are described in (Coquel et al., 2018).

### Western-blot

Cellular extracts were resuspended in Laemmli buffer (65.8 mM Tris, 26.3% glycerol, 2.1% SDS, and Bromophenol blue) and boiled at 95°C for 5 min. Proteins were separated by SDS-PAGE using home-made or precast gels (Bio-Rad) with suitable percentage then transferred on nitrocellulose membranes (GE Healthcare or Bio-Rad). Membranes were blocked with 5% non-fat milk in TBS-T (10 mM Tris pH 8.0, 150 mM NaCl, 0.5% Tween 20) for 1 hr then incubated with the primary antibodies overnight. Membranes were washed 3 times with TBS-T then incubated with the corresponding secondary antibody. Finally, membranes were developed with Clarity Western ECL Blotting Substrate (Bio-Rad) and images were acquired using a ChemiDoc System (Bio-Rad). Antibodies against the following proteins were used: Ser345 Phospho-Chk1 (Cell Signaling Technology 2348), Chk1 (Santa Cruz sc-8408), PCNA (Sigma-Aldrich P8825), Ser4/8 Phospho-RPA32 (Bethyl A300-245A), RPA32 (Calbiochem NA18), histone H3 (Abcam ab62642), GNL3 (Bethyl A300-600A, Santa Cruz sc-166460 or Sigma-Aldrich SAB1407312), Ser33 Phospho-RPA32 (Bethyl A300-246), Tubulin (Sigma Aldrich T5168), CDC45 (Santa Cruz sc-20685), Ser40 Phospho-MCM2 (Abcam ab133243), MRE11 (Novus NB100-142), BRCA1 (Santacruz sc-642), CtIP (Abcam ab70163), RAD51 (Santa Cruz sc-8349) and FLAG (Sigma Aldrich F1804).

### esiRNA screening

The 25 esiRNA (Sigma-Aldrich) corresponding to 24 candidates plus 1 negative control (EGFP) are described in SupTable1. HCT116 were seeded in 96 wells plates and transfected with esiRNAs using Oligofectamine (ThermoFisher). After 48 hours, transfected cells were subjected to 4 hrs treatment with 1 μM camptothecin then fixed for 15 min using 4% paraformaldehyde (PFA). Cells were permeabilized with 75% EtOH for 30 min on ice. 96 wells plate was incubated with primary antibody against Ser139 Phospho-H2A.X (Millipore 05-636) for 60 min then with secondary antibody anti-mouse coupled with Alexa568 (ThermoFisher A-11011) and finally with DAPI for 30 min. All the washes were performed with PBS-BSA 1%. 96 wells were scanned using a Nexcelom Celigo and images were analyzed using Celigo software. DAPI staining was used to measure the level of Ser139 Phospho-H2A.X in the nucleus for each esiRNA.

### Proximity Ligation Assay (PLA)

Cells were grown on coverslips to reach 70-80% confluency then fixed with 2% paraformaldehyde (PFA) and 0.02% sucrose in PBS for 20 min at room temperature. When specified cells were incubated with EdU (5-ethynyl-2’-deoxyuridine) for the indicated times. Cells were permeabilized with 0.5% Triton X100-PBS for 20 min then washed PBS-3% BSA. EdU was conjugated to biotin-TEG-azide (Eurogentec) using Click-it reaction (30 min at room temperature) using indicated concentrations (10 mM sodium Ascorbate, 5 μM biotin-TEG-azide, 3 μM CuSO_4_). For Click-it negative controls, biotin-TEG-azide was replaced by DMSO.

Coverslips were incubated with primary antibodies in PLA blocking solution (Sigma-Aldrich) overnight at 4°C then washed with PBS. PLA probes (anti-mouse minus DUO92004 and anti-rabbit plus DUO92002, Sigma-Aldrich) were incubated together in PLA blocking solution for 20 min then added on the coverslips for 1 h at 37°C then washed 2 times with buffer A (150 mM NaCl, 10 mM Tris, 0.5 % Tween). PLA kit was used (DUO92014, Sigma-Aldrich) for the following steps. Coverslips were incubated with ligase (1/40 dilution in ligase buffer) for 30 min at 37°C. Coverslips were washed 2 times with buffer A and incubated with polymerase (1/80 dilution in amplification buffer) for 100 min at 37°C. Coverslips were washed 2 times with buffer B (200 mM NaCl, 400 mM Tris-Base), dried and then mounted on glass slides with DAPI containing mounting medium (DUO82040 Sigma-Aldrich). Cells were analyzed by fluorescence microscopy and quantification the number of foci was performed using Fiji software. Antibodies against the following proteins were used: Biotin (Bethyl A150-109 and Jackson Immunoresearch 200-002-211), ORC2 (Bethyl A302-734A), CENP-A (Thermo Fisher MA1-20832), FLAG (Sigma Aldrich F1804) and GNL3 (Bethyl A300-600A and Santa Cruz sc-166460).

### Flow Cytometry

When indicated cells were first labeled with 20 μM IdU for 10 min and then fixed with ice­cold 70% ethanol. Then cells were treated with RNase during 60 min and then for 30 minutes with 2M HCl. Next, the cells were incubated with a BrdU/IdU antibody from BD Biosciences (347580) for 60 min or with an anti-pH3S10 (Cell Signaling 9701) overnight, and then with an Alexa 488 conjugated anti-mouse IgG (Invitrogen) at room temperature for 30 min. Finally, the cells were stained with 5 μg/ml of propidium iodide in PBS and analyzed using a MACSquant analyzer (Miltenyi Biotec). Results were analyzed using Flowjo (https://www.flowjo.com).

### Replication analysis by DNA Combing

Asynchronous cells were labeled 20 min with IdU, 20 min with CldU and then chased 90 min with thymidine. Purification of HMW gDNA, DNA combing and replication analysis was performed as in (Bialic et al., 2015) with the following modifications. Agarose plugs containing gDNA were washed in TNE50 containing 100 mM NaCl, digested O/N at 42°C with 3U β-agarase (New England Biolabs) and again for 2 hrs with 2U β-agarase. DNA was combed in MES buffer also containing 100 mM NaCl. Briefly, genomic DNA was combed on silanized coverslips, denatured with NaOH, and sites of DNA synthesis revealed using anti-IdU (red), anti-CldU (green), and anti-ssDNA (blue) antibody pairs. Primary antibodies were rat anti-BrdU (clone BU1/75, Abcam ab6326) for CldU, mouse anti-BrdU (clone B44, Becton Dickinson), for IdU and mouse autoanti-ssDNA (from DSHB) for DNA. Washes were performed with PBS-T containing 0.05% Triton X100. Secondary antibodies were Alexa488 Goat anti-rat IgG, Alexa546 Goat anti-mouse IgG, Alexa647 Goat anti-Mouse IgG2a (Life Technologies). Imaging was performed on a Zeiss AxioImager Z1 microscope with YFP, Cy3 and Cy5 filter blocks, equipped with a 40× objective (EC Plan Neofluar 1.3 NA oil) and scMOS ZYLA 4.2 MP camera (2048*2048 pixels, 6.5µm pixel size). Red-to-green signals show fork direction (yellow arrow). Fork velocity (FV) is calculated by dividing the length of the green tract by the pulse time (in kb/min). Global instant fork density (GIFD) was calculated using the formula that accounts for the doubling of DNA during S phase:

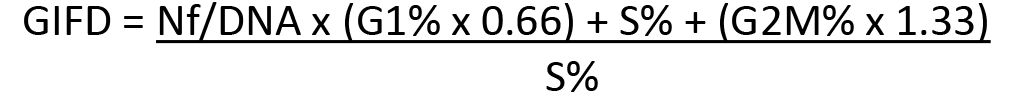

where Nf is the number of bicolor forks, DNA the total length of DNA measured (in Mb) and G1%, S% and G2M% the fraction of cells in G1, S and G2 or M phases, respectively, calculated from flow cytometry profiles using the same cells as for DNA combing.

### Isolation of proteins on Nascent DNA (iPOND)

iPOND was performed largely as previously described (Lossaint et al., 2013; Ribeyre et al., 2016). HeLa S3 cells were pulse labeled with 10 μM EdU for indicated times and chases were performed with 10 μM thymidine. Cells were fixed with 1% formaldehyde for 5 min or 2% for 15 min followed or not by quenching of formaldehyde by 5 min incubation with 0.125 M glycine. Fixed samples were collected by centrifugation at 1000 g for 3 min, washed three times with PBS and stored at -80^⁰^C. Cells were permeabilized with 0.5% triton for 30 min and click chemistry was used to conjugate biotin-TEG-azide (Eurogentec) to EdU-labelled DNA in PBS containing 10 mM sodium Ascorbate, 10 μM biotin-TEG-azide, 2 mM CuSO_4_. Cells were re-suspended in lysis buffer (10 mM Hepes-NaOH; 100 mM NaCl; 2 mM EDTA PH8; 1 mM EGTA; 1 mM PMSF; 0.2% SDS; 0.1% Sarkozyl) and sonication was performed using a Qsonica sonicator with the following settings: 30% power, 20 sec constant pulse and 50 sec pause for a total sonication time of 5 min on ice with water. Lysates were centrifuged at 15,000 g for 10 min at room temperature. Supernatants were normalized by DNA quantification using a nanodrop device. Biotin conjugated DNA-protein complexes were captured using overnight incubation with magnetic beads coated with streptavidin (Ademtech). Captured complexes were washed with lysis buffer and 500 mM NaCl. Proteins associated with nascent DNA were eluted under reducing conditions by boiling into SDS sample buffer for 30 min at 95°C and analyzed by Western-blot.

### DNA fibers labelling

DNA fibers labelling was performed as previously described (Lossaint et al., 2013; Ribeyre et al., 2016). Cells were labeled with 25 μM IdU, washed with warm media and exposed to 50 μM CldU. Cells were lysed and DNA fibers were stretched onto glass slides are left to air dry then are fixed in methanol/acetic acid (3:1) for 10 min. The DNA fibers were denatured with 2.5 M HCl for 60 min, washed with PBS and blocked with 2% BSA in PBS-Tween for 60 min. IdU replication tracks were revealed with a mouse anti-BrdU/IdU antibody from BD Biosciences (347580) and CldU tracks with a rat anti-BrdU/CldU antibody from Eurobio (ABC117-7513). The following secondary antibodies were used: Alexa fluor 488 anti-mouse antibody (Life A21241) and Cy3 anti-rat antibody (Jackson Immunoresearch 712-166-153). Fibers were visualized and imaged by Carl Zeiss Axio Imager Apotome using 40X Plan Apo 1.4 NA oil immersion objective. Replication tracks lengths were analyzed using ImageJ software. Statistical analysis was performed using Graphpad Prism software.

### Immunofluorescence

Cells were grown on coverslips to reach 70-80% confluency then directly with 4% paraformaldehyde (PFA) in PBS for 20 min at room temperature. When indicated, incubation with cytoskeletal (CSK) buffer (10 mM PIPES pH 6.8, NaCl 100mM, sucrose 300 mM, MgCl_2_ 3mM, EGTA 1mM, Triton X-100 0.5%) was performed. Cells were permeabilized by with 0.2% Triton X100-PBS for 10 min then transferred into 0.1% Tween-PBS for 5 min. Coverslips were then incubated with primary antibodies in 0.1% Tween-5% BSA-PBS for 1-2 hrs, washed with 0.1% Tween-PBS, then incubated with secondary antibodies in Tween 0.1%-BSA 5%-PBS for 1 hr. All the incubations were carried out in darkness in a humidified chamber at room temperature. Finally, coverslips are washed again with 0.1% Tween-PBS, incubated with Hoechst to label DNA for 5 min, and then mounted on glass slides with Prolong (Life). Cells were analyzed by fluorescence microscopy. Antibodies against the following proteins were used: FLAG (Sigma Aldrich F1804), Streptavidin-Alexa Fluor 488 (Life S32354), NOP1 (Novus NBP2-46881) and GNL3 (Bethyl A300-600A).

### Replication timing experiments and microarrays

Cells were incubated with 50 µM of BrdU for 90 min and collected, washed three times with PBS and then fixed in ethanol 75%. Cells were re-suspended in PBS with RNAse (0.5 mg/ml) and then with propidium iodide (50 µg/ml) followed by incubation in the dark at room temperature for 30 min with low agitation. Two fractions of 150,000 cells, S1 and S2 corresponding to Early and Late S-phase fractions respectively, were sorted by flow cytometry using a Becton Dickinson FACS Melody. Whole DNA was extracted with lysis buffer (50 mM Tris pH 8, 10 mM EDTA, 300 mM NaCl, 0.5% SDS) and 0.2 mg/ml of Proteinase K for 2 hrs at 65°C. Neo-synthesized DNA were immunoprecipitated with BrdU antibodies (Anti-BrdU Pure, BD Biosciences, #347580) as previously described (Fernandez-Vidal et al., 2014). To control the quality of enrichment of early and late fractions in S1 and S2, qPCR was performed with BMP1 oligonucleotides (early control) and with Dppa2 oligonucleotides (late control; data not shown, (Hiratani et al., 2008)). Microarray hybridization requires a minimum of 1000 ng of DNA. To obtain sufficient specific immunoprecipitated DNA for this hybridization step, whole genome amplification was conducted (WGA, Sigma) on immunoprecipitated DNA. A post WGA qPCR was performed to preserve specific enrichment in both S1 and S2 fractions. Early and late amplified neo-synthesized DNA were then labeled with Cy3 and Cy5 ULS molecules, respectively (Genomic DNA labeling Kit, Agilent). The hybridization was performed according to the manufacturer instructions on 4×180K mouse microarrays (SurePrint G3 Mouse CGH Microarray Kit, 4x180K, AGILENT Technologies, reference genome: mm9). Microarrays were scanned with an Agilent High-Resolution C Scanner using a resolution of 3 µm and the autofocus option. Feature extraction was performed with the Feature Extraction 9.1 software (Agilent Technologies). For each experiment, the raw data sets were automatically normalized by the Feature extraction software. Analysis was performed using the STAR-R software described in (Hadjadj et al., 2020). The statistical comparison was conducted between early and late domains from both cell lines in order to determine segments where replication timing changes. Graphical representation was generated with START-R suit.

### Chromatin immunoprecipitation and deep sequencing (ChIP-seq)

About 20.10^6^ of Hela S3 cells per sample were prepared for sonication following the True-ChIP chromatin shearing kit protocol for High Cell concentration from Covaris. Cells were cross-linked in 1% methanol-free formaldehyde during 5 min before cell lysis and nuclei preparation. Washed nuclei were sonicated for 15 min at 6°C to obtain DNA fragments of 100-800pb using the E220evolution Covaris machine following parameters indicated in the provided protocol. After dilution with one volume of immunoprecipitation dilution buffer (Covaris), sonicated samples were pre-cleared with 3 µL/mL of protein G magnetic beads (Ademtech) during 1 hr at 4°C. Each sample was then normalized to an equal amount of protein (associated to pre-cleared chromatin) and input samples were collected after this step. Normalized samples were then incubated with 1 µg of GNL3 antibody (Bethyl A300-600A) overnight at 4°C, before incubation with 20 µL/mL of protein G magnetic beads (previously blocked overnight at 4°C in immunoprecipitation dilution buffer with 1% BSA) during 4 hrs at 4°C. Chromatin bound to beads was then washed 5 min at room temperature in each following buffers: low salt buffer (150 mM NaCl, 20 mM Tris HCl pH=8, 2 mM EDTA, 1% Triton, 0.1% SDS); high salt buffer (500 mM NaCl, 20 mM Tris HCl pH=8, 2 mM EDTA, 1% Triton, 0.1% SDS); LiCl buffer (0.25 M LiCl, 10 mM Tris-HCl pH=8, 1 mM EDTA, 1% Sodium deoxycholate, 1% NP-40); TE buffer (10 mM Tris-HCl pH=8, 1 mM EDTA). Washed beads were eluted in 200 µL of elution buffer (100 mM NaHCO3, 1% SDS) during 15 min at 30°C with shaking. Eluted chromatin and input samples were reverse-crosslinked overnight at 65°C with 0.2 M NaCl and 0.02 mg/mL of RNAse A and incubated 1 hr with Proteinase K (400 µg/mL final concentration). DNA was purified using the ChIP DNA Prep Adem kit (Ademtech) following the provided protocol. DNA bound to beads was eluted in 50 µL of elution buffer. Quantity of DNA was measured with the Qubit 1X dsDNA HS Assay kit (Invitrogen), using a Qubit 2.0 fluorometer (Thermofisher scientific). GNL3 ChIP was repeated three times and 10 ng of each ChIP and each corresponding input were pooled together and send to the MGX sequencing platform of Montpellier, France (https://www.mgx.cnrs.fr/). DNA banks were sequenced using the Illumina-Novaseq-6000 machine to obtain 150 bp paired-end reads. Sequencing data were processed and analyzed using the online Galaxy platform (https://usegalaxy.org/). Reads were aligned on the February 2009 human reference genome (GRCh37/Hg19) using Bowtie2 tool with default parameters. GNL3 Peaks were discovered using MACS2 callpeak tool using input as control file with a q-value<0.005. ORC2 peaks file was taken from Miotto et al. (Miotto et al., 2016).

### Chromatin Fractionation

Cells were seeded at 80% confluency and collected by trypsinization followed by centrifugation for 3 min (1200g) at room temperature. The pellets were washed with PBS then resuspended with CSK buffer (10 mM PIPES pH 6.8, 100 mM NaCl, 300 mM Sucrose, 1 mM MgCl_2_, 1 mM EGTA, 0.5 mM DTT, 0.1% Triton X-100, 1 mM ATP, 1X protease inhibitor) and kept for 10 min on ice. Lysed cells were then centrifuged for 3 min (3000g) at 4°C. The resulting supernatant presenting the soluble protein fraction was transferred to another Eppendorf tube and the pellet was washed with CSK buffer for 10 min on ice followed by centrifugation for 3 min (3000g) at 4°C. the resulting pellet which represents the in-soluble fraction of proteins was then resuspended in 2X Laemmli buffer and incubated at 95°C for 10 min before western blot analysis.

### Bio-ID

Flp-In T-Rex 293 cell lines were stably transfected with Flag-BirA-GNL3. Cells seeded at 75% confluency were incubated with 10 µg/ml of doxycycline for 16 hrs and then with 50 µM biotin for 4 hrs. Cells were washed once with PBS and lysed with RIPA/SDS buffer (50 mM Tris-HCl pH 7.5, 150 mM NaCl, 1 mM EDTA, 1 mM EGTA, 1% NP-40, 0.2% SDS, 0.5% Sodium deoxycholate) complemented with 1X complete protease inhibitor and 250U benzonase (Sigma-Aldrich, CE1014). Lysed cells were incubated on a rotating wheel for 1 hr at 4°C followed by sonication on ice with 30% amplitude for 3 cycles of 10 sec (2 sec ON-2sec resting) separated with 10 sec of resting. Sonicated lysate was next centrifuged for 30 min (7750g) at 4°C, the cleared supernatant was transferred to a new tube and protein concentration was quantified using Bradford protein assay. For each condition, 500 µg of proteins were incubated with 30 µl of Streptavidin-Agarose beads (Sigma-Aldrich, CS1638) on a rotating wheel for 3 hrs at 4°C. Beads were next washed sequentially with 1 ml of each buffer starting with lysis buffer, wash buffer 1 (2% SDS in H_2_O), wash buffer 2 (0.2% sodium deoxycholate, 1% Triton X-100, 500 mM NaCl, 1mM EDTA, and 50mM Hepes pH 7.5), wash buffer 3 (250 mM LiCl, 0.5% NP-40, 0.5% sodium deoxycholate, 1mM EDTA, 500mM NaCl and 10mM Tris pH 8) and finally wash buffer 4 (50 mM Tris pH 7.5 and 50 mM NaCl). Biotinylated proteins were eluted from the magnetic beads using 40 µl of 2X Laemmli buffer and incubated at 95°C for 10 min.

### Proteomics analysis of Bio-ID samples

Biotinylated proteins were migrated on SDS PAGE for a short migration. After reduction (DTT 1 M, 30 min at 60°C) an alkylation (IAA 0.5 M, 30 min RT) proteins were digested using trypsin (Gold, Promega, 1ug / sample, overnight at 37°C). For LC MSMS analysis, samples were loaded onto a 50 cm reversed-phase column (75 mm inner diameter; Acclaim PepMap 100 C18; Thermo Fisher Scientific) and separated with an UltiMate 3000 RSLC system (Thermo Fisher Scientific) coupled to a QExactive HF system (Thermo Fisher Scientific). Separation of the peptides was performed following a gradient from 2 to 25% buffer B (0.1% AF in 80% ACN) for 100 min at a flow rate 300 nl / min, then 25 to 40% in 20 min and finally 40 to 90% in 3 minutes. Tandem mass spectrometry analyses were performed in a data-dependent mode. Full scans (350–1,500 m/z) were acquired in the Orbitrap mass analyzer with a resolution of 60,000 at 200 m/z. For MS scans, 3e6 ions were accumulated within a maximum injection time of 60 ms. The 12 most intense ions with charge states ≥2 were sequentially isolated (1e5) with a maximum injection time of 100 ms and fragmented by higher-energy collisional dissociation (normalized collision energy of 28) and detected in the Orbitrap analyzer at a resolution of 30,000. Raw spectra were processed with MaxQuant v 1.6.5.0 (Cox and Mann, 2008) using standard parameters with match between runs option. Spectra were matched against the UniProt reference proteome (release 2019_06; http://www.uniprot.org) of Homo sapiens and 250 frequently observed contaminants, as well as reversed sequences of all entries. The maximum false discovery rate for peptides and proteins was set to 0.01. Representative protein ID in each protein group was automatically selected using the in-house developed Leading tool (Raynaud et al., 2018).

### Immunoprecipitation

Whole-cell extracts of K562 cells were prepared using lysis buffer (50 mM Tris-HCl pH 8, 150 mM NaCl, 5 mM EDTA pH8, 0.5% NP40) supplemented with protease inhibitor cocktail (Roche), 1mM PMSF, 1mM MgCl_2_ and Benzonase Nuclease 250 units/ 10 millions of cells (E1014-25KU, Sigma). Immunoprecipitations were performed overnight at 4°C with protein G Dynabeads (Thermo Fisher Scientific) coupled to either rabbit immunoglobulin G (IgG) (P120-201, Bethyl Laboratories) or rabbit ORC2 antibody (A302-734A, Bethyl Laboratories). Beads were washed 4 times with lysis buffer, then washed 3 times with 50 mM Tris HCl pH8. The immunoprecipitated complexes were eluted in 50 mM Tris HCl pH8 containing 1% SDS for 15 min à 56°C with agitation. IP samples were mixed with 1X Bolt Sample Reducing agent (Thermo Fisher Scientific) and 1X Bolt LDS Sample Buffer (Thermo Fisher Scientific), loaded and resolved on pre-cast Bolt Bis-Tris gels (Thermo Fisher Scientific), then transferred onto nitrocellulose membrane (GE Healthcare). Membranes were blocked in 5% fat-free milk in PBS, incubated overnight at 4°C with primary antibodies directed against ORC2 (A302-734A, Bethyl Laboratories) and GNL3 (sc-166460, Santa Cruz Biotechnology). A cognate secondary antibody coupled to horseradish peroxidase was used and revealed with the Super Signal West Dura Extended Duration Substrate kit (Thermo Fisher Scientific). Acquisition was performed using the Fusion FX (Vilber) and image analysis was performed using ImageJ (https://imagej.nih.gov/ij/).

### Proteomics analysis of immunoprecipitation

Sample preparation: Tryptic peptides from the immunoprecipitated complexes (=eluate) were obtained by Strap Micro Spin Column according to the manufacturer’s protocol (Protifi, NY, USA). Briefly: proteins from 140 µL of the eluate were diluted 1:1 with 2x reducing-alkylating buffer (20 mM TCEP, 100 mM Chloroacetamide in 400 mM TEAB pH 8.5 and 4% SDS) and left 5 min at 95°C to allow reduction and alkylation in one step. Strap binding buffer was applied to precipitate proteins on quartz and proteolysis took place during 14 hrs at 37°C with 1 µg Trypsin sequencing grade (Promega). After speed-vacuum drying of eluted peptides, these were solubilized in 0.1% trifluoroacetic acid (TFA) in 10% Acetonitrile (ACN). Liquid Chromatography-coupled Mass spectrometry analysis (LC-MS): LC-MS analyses were performed on a Dionex U3000 HPLC nanoflow system coupled to a TIMS-TOF Pro mass spectrometer (Bruker Daltonik GmbH, Bremen, Germany). One μl was loaded, concentrated and washed for 3 min on a C18 reverse phase precolumn (3 μm particle size, 100 Å pore size, 75 μm inner diameter, 2 cm length, from Thermo Fisher Scientific). Peptides were separated on an Aurora C18 reverse phase resin (1.6 μm particle size, 100Å pore size, 75 μm inner diameter, 25 cm length mounted onto the Captive nanoSpray Ionization module, (IonOpticks, Middle Camberwell Australia) with a 60 minutes overall run-time gradient ranging from 99% of solvent A containing 0.1% formic acid in milliQ-grade H2O to 40% of solvent B containing 80% acetonitrile, 0.085% formic acid in mQH2O. The mass spectrometer acquired data throughout the elution process and operated in DDA PASEF mode with a 1.1 second/cycle, with Timed Ion Mobility Spectrometry (TIMS) mode enabled and a data-dependent scheme with full MS scans in PASEF mode. This enabled a recurrent loop analysis of a maximum of the 120 most intense nLC-eluting peptides which were CID-fragmented between each full scan every 1.1 second. Ion accumulation and ramp time in the dual TIMS analyzer were set to 50 ms each and the ion mobility range was set from 1/K0 = 0.6 Vs cm-2 to 1.6 Vs cm-2. Precursor ions for MS/MS analysis were isolated in positive mode with the PASEF mode set to « on » in the 100-1.700 m/z range by synchronizing quadrupole switching events with the precursor elution profile from the TIMS device. The cycle duty time was set to 100%, accommodating as many MSMS in the PASEF frame as possible. Singly charged precursor ions were excluded from the TIMS stage by tuning the TIMS using the otof control software, (Bruker Daltonik GmbH). Precursors for MS/MS were picked from an intensity threshold of 2.500 arbitrary units (a.u.) and resequenced until reaching a ‘target value’ of 20.000 a.u taking into account a dynamic exclusion of 0.40 s elution gap. Protein quantification and comparison: The mass spectrometry data were analyzed using Mascot version 2.5.1 (http://www.matrixscience.com/). The database used was a concatenation of Homo sapiens sequences from the Swissprot databases (release June 2020: 563,972 sequences; 203,185,243 residues) and an in-house list of frequently found contaminant protein sequences. The enzyme specificity was trypsin’s. The precursor and fragment mass tolerances were set to 20ppm. Oxidation of methionines was set as variable modifications while carbamidomethylation of cysteines was considered complete. False discovery rate (FDR) was kept below 1% on both peptides and proteins. For comparative analysis, peptide count results from Mascot were assembled with the MyPROMS (Poullet et al., 2007) software (version 3.1).

## Expanded View Figures

**Expanded View Figure 1. A.** Scheme explaining the mini esiRNA screen to determine the best candidates for further characterization. HCT116 cells were transfected in 96 wells plate with each esiRNA from the library (SupTable1). After 48 hrs cells were treated with 1 μM camptothecin for 4 hrs and subjected to immunofluorescence using an antibody directed against γH2A.X. The level of γH2A.X within nuclei was analyzed using a Celigo high-throughput microscope. **B.** Candidates were sorted according to their level of γH2A.X upon camptothecin treatment upon 5 independent experiments. **C.** Graphic representation of CldU/IdU ratio in response to CPT in HeLa S3 cells in 3 independent experiments. **D.** Western-blot analysis of HeLa S3 cells treated with 1 μM camptothecin (CPT) during the indicated time. **E.** Western-blot analysis of HeLa S3 cells treated with 10 μM etoposide (ETP) during the indicated time. **F.** Graphic representation of the average level of resection (indicated in red) in response to HU in HeLa S3 cells in 3 independent experiments. **G.** HeLa S3 cells were sequentially labelled for 30 min with IdU and for 30 min with CldU then treated with 1 μM CPT for 240 min. The ratio between CldU and IdU is plotted, the red line indicates the median. For statistical analysis Mann-Whitney test was used; ****p<0.0001. The average level of resection (indicated in red) in 3 independent experiments is also shown. **H.** HeLa S3 cells were sequentially labelled for 30 min with IdU and for 30 min with CldU then treated with 10 μM ETP for 120 min. The ratio between CldU and IdU is plotted, the red line indicates the median. For statistical analysis Mann-Whitney test was used; ****p<0.0001. The average level of resection (indicated in red) in 3 independent experiments is also shown. **I.** Western-blot analysis of cells depleted for GNL3, MRE11 or CtIP. **J.** Average level of resection in response to HU (indicated in red) in 3 independent experiments. **K.** Average level of resection in response to HU (indicated in red) in 3 independent experiments upon complementation with exogenous GNL3-WT.

**Expanded View Figure 2. A.** Flow cytometry experiment of HeLa S3 cells. Nascent DNA was labelled with IdU and total DNA stained with propidium iodide. **B.** Percentage of HeLa S3 cells positive for histone H3 phosphorylated on Histone 10 (pH3S10) upon thymidine block and release. The error bars represent the average of 3 independent experiments. **C.** Graphic representation of the 3 independent DNA combing experiments. The average value is indicated in red. **D.** Loss of RIF1 has effect on replication timing in specific genomic loci. Cells were pulse-labelled with BrdU for 90 min and sorted by flow cytometry in two fractions, S1 and S2, corresponding to early and late S-phase. Neo-synthesized DNA was immunoprecipitated with BrdU antibodies. Early and late neo-synthesized DNAs were labeled with Cy3 and Cy5 and hybridized on microarrays. After processing analyzing with the START-R software, replication-timing profiles can be obtained from two replicates. Shown are the zoomed microarray profiles of the timing of replication on chromosome 1 and chromosome 15 as example. Blue lines represent replication timing from siControl cells and red lines represent siRIF1 cells and grey spots represent the log ratio intensity for each probe of the microarray. Significantly disturbed regions are detected by START-R software and advanced regions are indicated with green line and delayed regions by a pink line.

**Expanded View Figure 3. A.** Western-blot analysis upon treatment with HU and inhibition of CDC7, WEE1 or ATR. **B.** Average level of resection in response to HU (indicated in red) in 3 independent experiments upon inhibition of ATR and CDC7. **C.** Average level of resection in response to HU (indicated in red) in 3 independent experiments upon inhibition of WEE1 and CDC7. **D.** Average level of resection in response to HU (indicated in red) in 3 independent experiments upon GNL3 depletion and CDC7 inhibition. **E.** Quantification of RPA phosphorylation (S4/8) upon GNL3 depletion and CDC7 inhibition. The error-bars represent the variation of 3 independent experiments. **F.** Western-blot analysis of HeLa S3 cells depleted or not for BRCA1 upon treatment with CDC7 inhibitor. **G.** Average level of resection in response to HU (indicated in red) in 3 independent experiments upon BRCA1 depletion and CDC7 inhibition.

**Expanded View Figure 4. A.** Immunofluorescence experiment in Flp-In T-Rex HEK293 cells showing the expression of GNL3-BirA-FLAG and the biotinylation by BirA (revealed by streptavidin coupled with Alexa-488) upon addition of biotin. **B.** Common hits found by mass spectrometry between ORC2 immunoprecipitation and GNL3 BioID. The localization was determined using The Human Protein Atlas database (https://www.proteinatlas.org/). **C.** Examples of GNL3 peaks on chromosome 19 obtained by ChIP-seq of GNL3, INPUT is shown as negative control. The ORC2 ChIP-seq data are obtained from (Miotto et al., 2016). **D.** PLA (proximity ligation assay) analyzing the proximity between ORC2 and GNL3 in HeLa S3 cells. **E.** PLA (proximity ligation assay) analyzing the proximity between GNL3 and CENP-A in HeLa S3 cells.

**Expanded View Figure 5. A.** PLA (proximity ligation assay) analyzing the proximity between ORC2 and GNL3-WT-FLAG or GNL3-dB-FLAG in HeLa Flp-In cells upon doxycycline induction using the indicated antibodies. **B.** PLA (proximity ligation assay) analyzing the proximity between ORC2 and GNL3-dB-FLAG in HeLa Flp-In cells upon doxycycline induction using the indicated antibodies. **C.** Average level of resection in response to HU (indicated in red) in 3 independent experiments upon expression of GNL3-WT or GNL3-dB. **D.** Western-blot analysis of Flp-In T-Rex HeLa cells expressing exogenous GNL3-WT or GNL3-dB mutants. Cells were transfected with siControl or siGNL3 for 48 hrs then expression of GNL3-WT and GNL3-dB (resistant to the siRNA against GNL3) was induced using the indicated doses of doxycycline for 16 hrs. **E.** Immunofluorescence analysis of Flp-in T-Rex HeLa cells expressing GNL3-dB at the indicated doses of doxycycline. **F.** Average level of resection in response to HU (indicated in red) in 3 independent experiments.

