## Supplementary figures and images for "The nucleolar protein GNL3 prevents resection of stalled replication forks"

### Expanded View Figure 1

#### Expanded View Figure 1

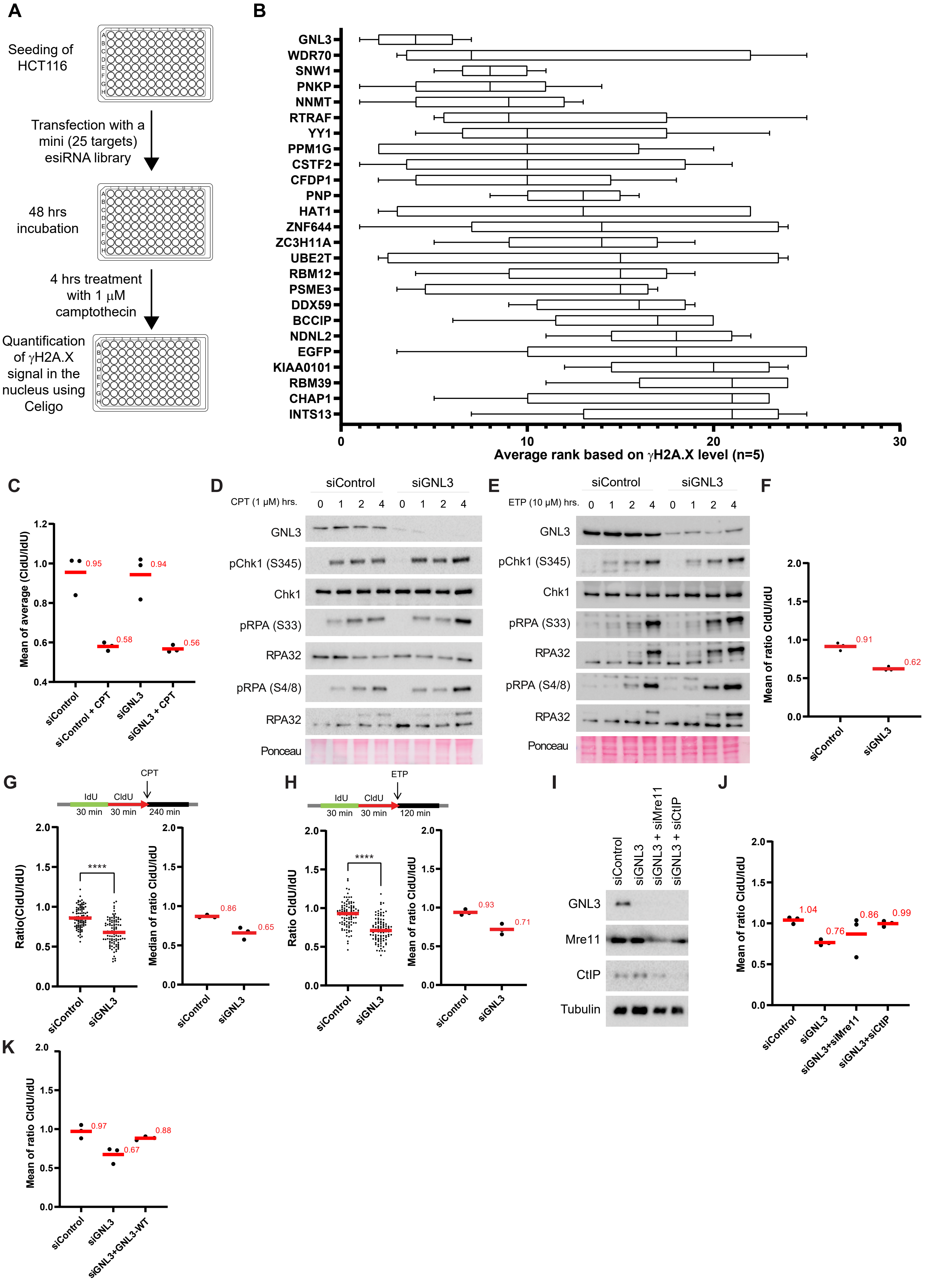

### Expanded View Figure 2

#### Expanded View Figure 2

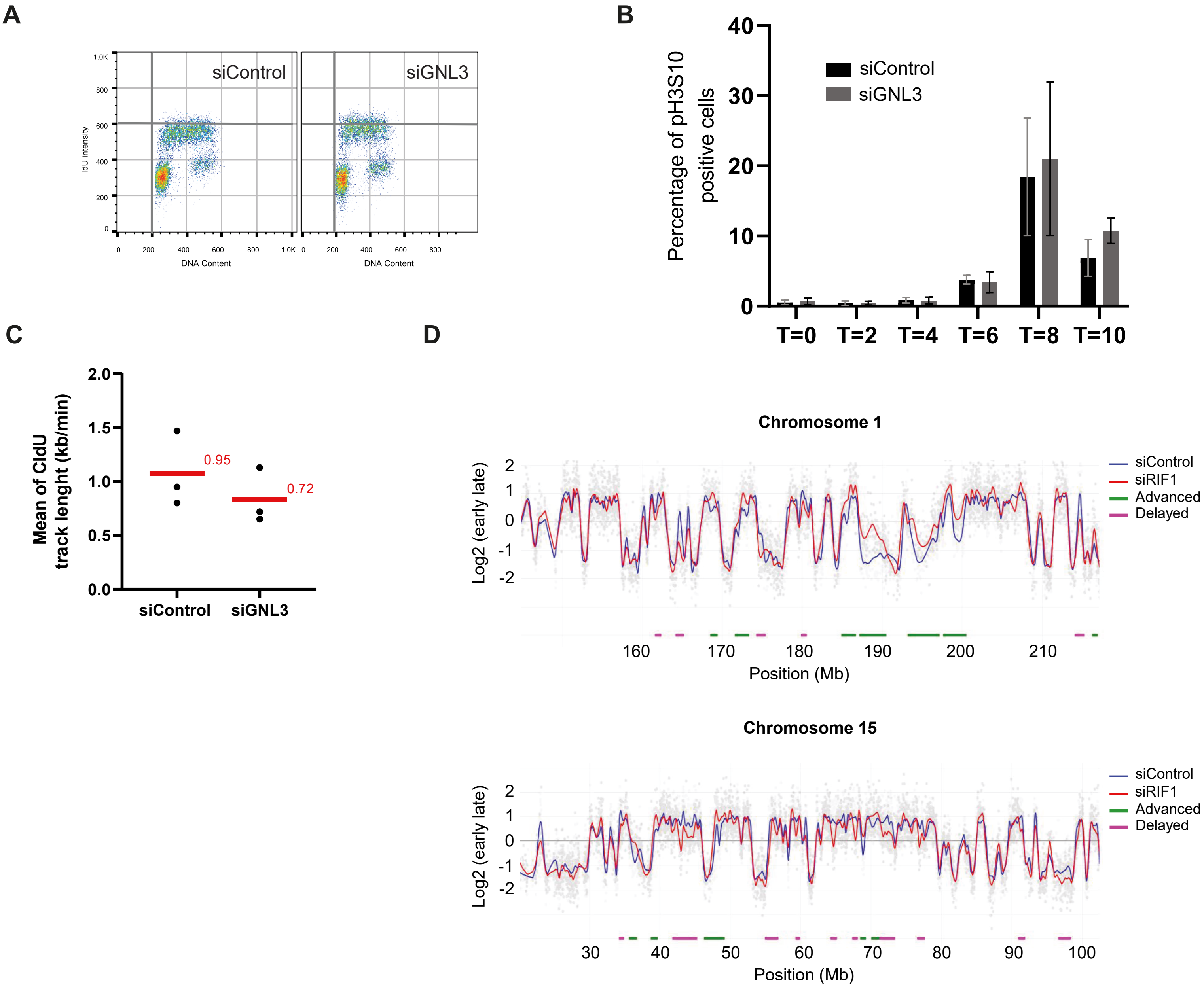

### Expanded View Figure 3

#### Expanded View Figure 3

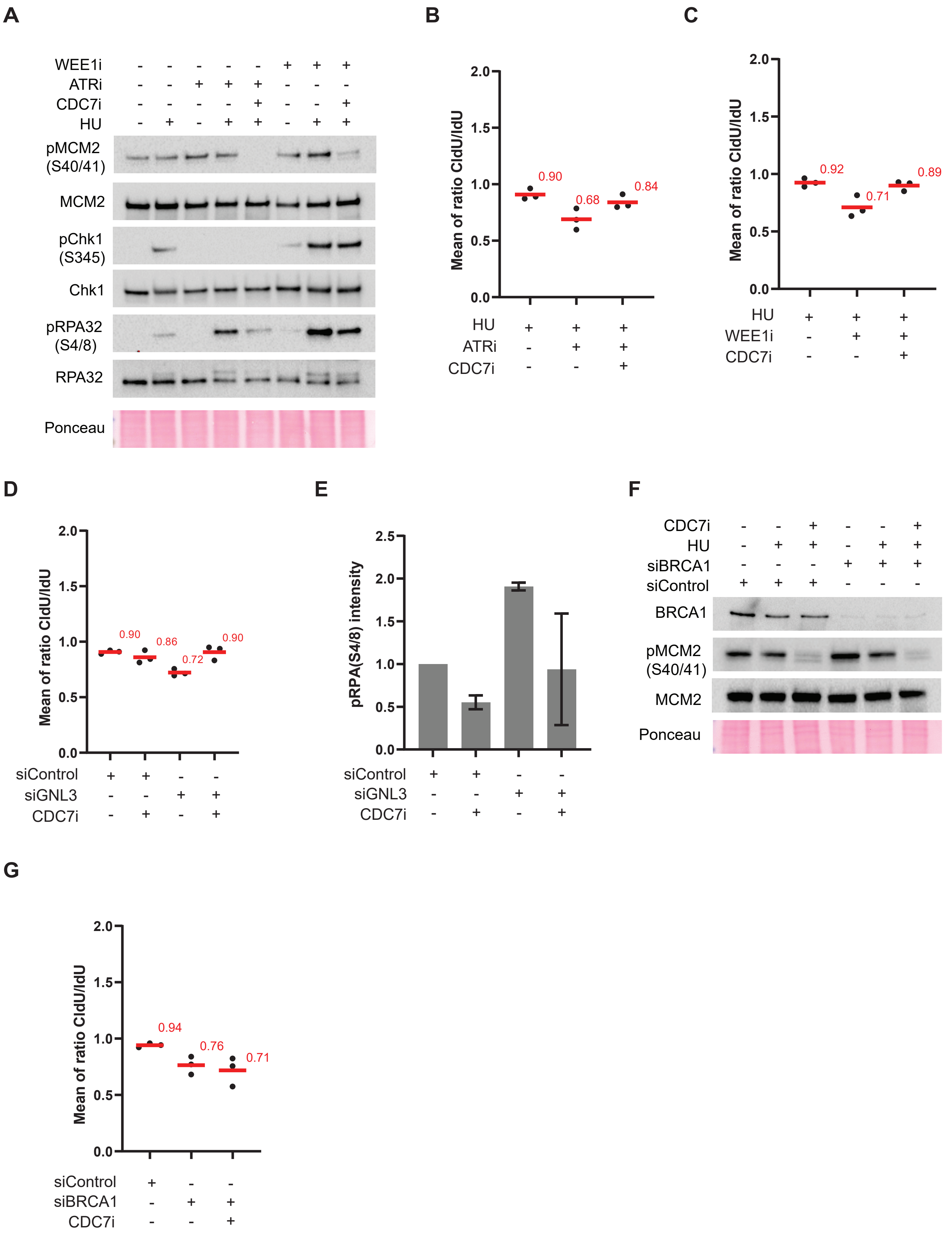

### Expanded View Figure 5

#### Expanded View Figure 5.

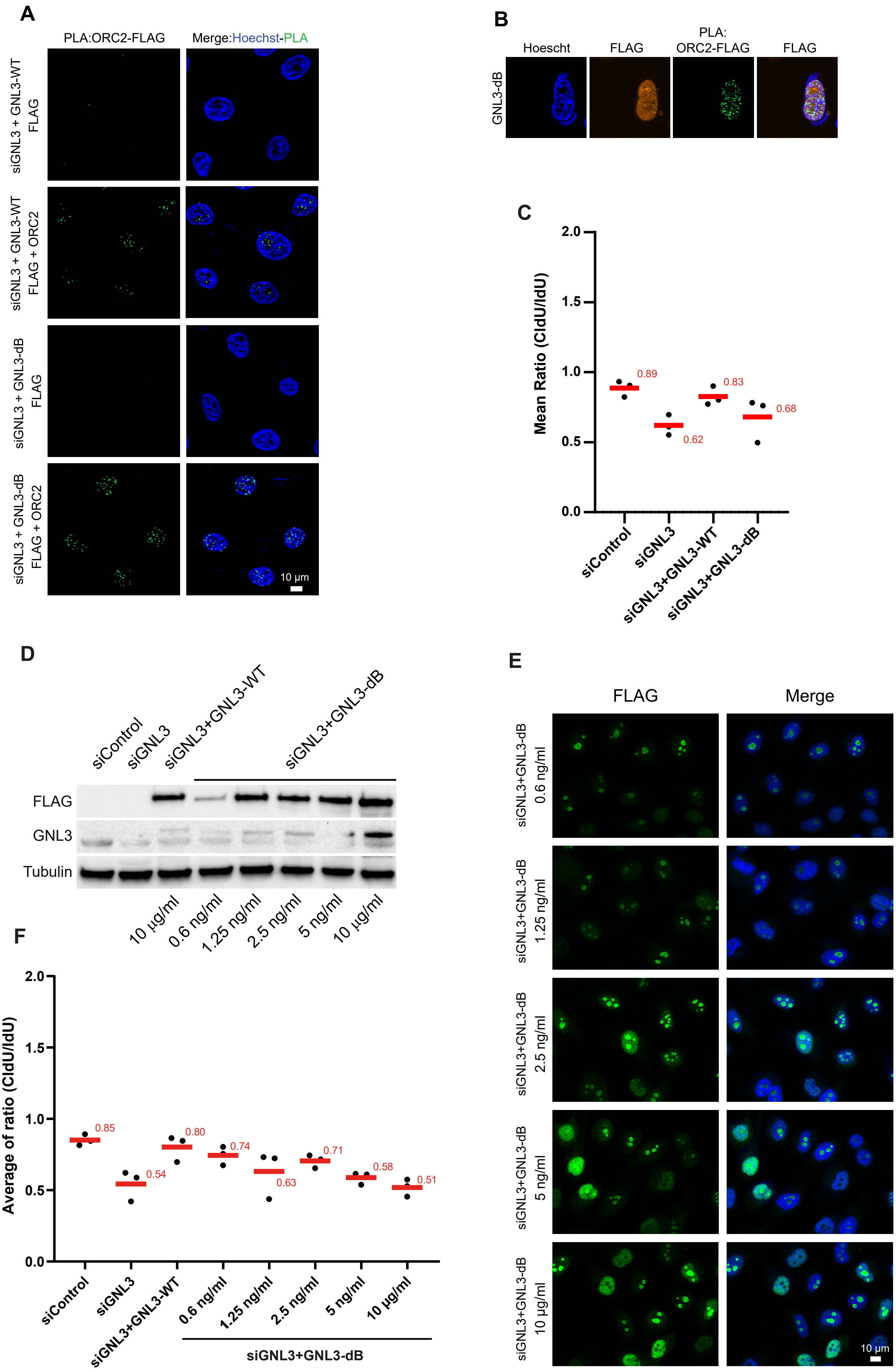
