## Supplementary material for "The nucleolar protein GNL3 prevents resection of stalled replication forks": Expanded View Figure 4

A

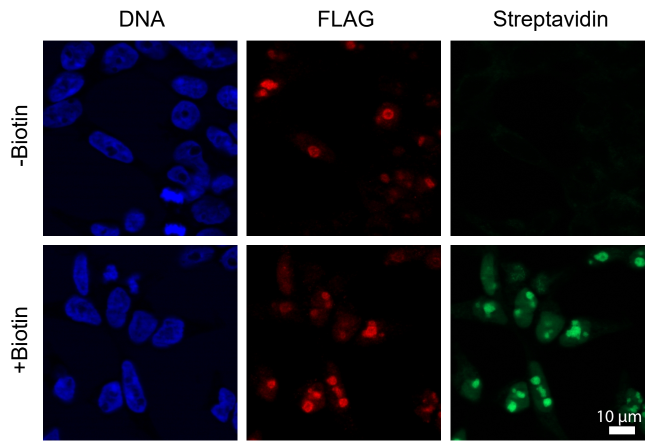


B

**Commons hits between ORC2-IP and GNL3 BioID**

| Present in the nucleolus | Not detected in the nucleolus |
|---|---|
| COIL | DDX49 |
| DDX10 | JMJD1C |
| DDX18 | MRE11 |
| DDX21 | PRPF3 |
| EBNA1BP2 | SF3B1 |
| ESF1 | SNRPA1 |
| EXOSC10 | THRAP3 |
| FTSJ3 | TMPO |
| GNL2 | TRAP1 |
| MKI67 | XRCC6 |
| MYBBP1A | ZC3H11A |
| NAT10 | |
| NCL | |
| NKRF | |
| NOP53 | |
| NPM1 | |
| PES1 | |
| RPF2 | |
| RPL5 | |
| TCOF1 | |
| TRMT1L | |
| UTP14A | |
| WDR36 | |
| XRN2 | |

C

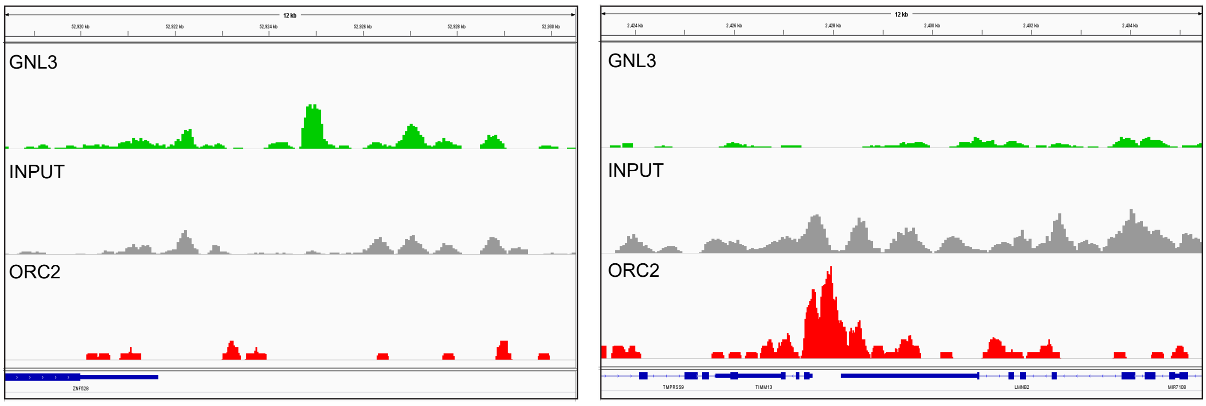


D

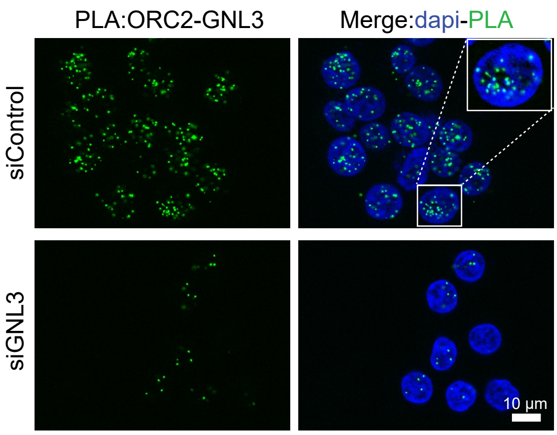


E

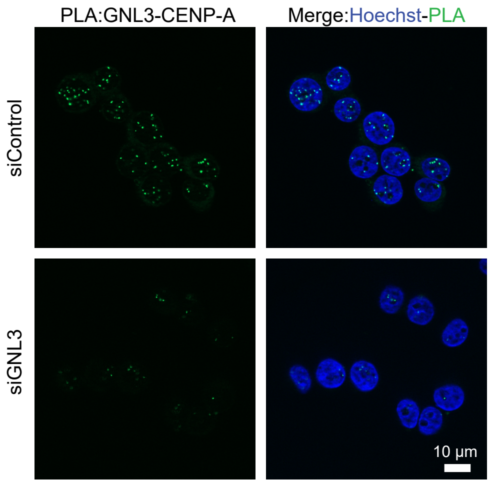
